# The cryo-EM structure of the bacterial type I DNA segregation ATPase filament reveals its conformational plasticity upon DNA binding

**DOI:** 10.1101/2021.03.22.436490

**Authors:** AV Parker, D Mann, SB Tzokov, LC Hwang, JRC Bergeron

## Abstract

The efficient segregation of replicated genetic material is an essential step for cell division. In eukaryotic cells, sister chromatids are separated via the mitotic spindles. In contrast, bacterial cells use several evolutionarily-distinct genome segregation systems. The most common of these is the Type I Par system. It consists of an adapter protein, ParB, that binds to the DNA cargo via interaction with the parS DNA sequence; and an ATPase, ParA, that binds nonspecific DNA and mediates cargo transport. However, the molecular details of how this system functions are not well understood.

Here, we report the cryo-EM structure of a ParA filament bound to its DNA template, using the chromosome 2 (Chr2) of *Vibrio cholerae* as a model system. We also report the crystal structures of this protein in various nucleotide states, which collectively offer insight into its conformational changes from dimerization through to DNA binding and filament assembly. Specifically, we show that the ParA dimer is stabilized by nucleotide binding, and forms a left-handed filament using DNA as a scaffold. Our structural analyses also reveal dramatic structural rearrangements upon DNA binding and filament assembly. Finally, we show that filament formation is controlled by nucleotide hydrolysis. Collectively, our data provide the structural basis for ParA’s cooperative binding to DNA and the formation of high ParA density regions on the nucleoid, and suggest a role for its filament formation.

## Introduction

DNA replication and segregation are essential in all life forms. In bacteria, partitioning (Par) systems are responsible for the efficient segregation of replicated genetic material (chromosome, low-copy-number plasmids) to daughter cells^1–4^. The *Par* loci are divided into three different types, classified by their motor protein: Type I *par* loci encode a P loop ATPase, ParA, which possesses a deviant Walker-type motif; the Type II ATPase, ParM, is actin-like; and the Type III GTPase, TubZ, is tubulin-like. The mechanisms of type II and III systems have been extensively characterized^4,5^. In contrast, the mechanism of the Type I segregation system remains elusive, with in particular discrepancy regarding ParA’s action during segregation^6^, despite being found in ~ 70% of bacteria.

The Type I segregation system locus encodes the ATPase ParA; an adapter protein, ParB; and centromere-like *parS* site(s). ParB binds specifically to its cognate *parS* site^7^, and was shown recently to have CTPase activity^8–10^, although the role of this activity remains unclear. ParA also binds to DNA in the presence of nucleotide, but unlike ParB this interaction is sequence-independent. Biochemical studies have shown that ParB stimulates ParA’s ATPase activity, promoting its dissociation from DNA^11–14^.

Several mechanistic models have been proposed for the Type I segregation system. A mitotic-like filament model was initially suggested^15–18^, similarly to type II and type III segregation systems. According to this model, ParA form filaments in the presence of ATP, and filament dissociation upon interaction with ParB causes a pulling of the chromosome. However, a range of evidence, including the lack of observation of ParA filaments *in vivo*, has led to questioning of this model^19^.

More recently, an alternative “diffusion ratchet” model has been proposed^20^, whereby the increase in in ATPase activity of ParA upon binding to ParB causes its dissociation from the DNA and an uneven distribution of ParA across the nucleoid; The partition complex then chases the ParA concentration gradient across the nucleoid whilst under confinement by the inner membrane, preventing diffusion of the partition complex into the cytosol. This model is supported by recent evidence using reconstituted systems and single-molecule measurements^13,21,22^; however, the molecular details of how it allows the diffusion of entire DNA molecules across the bacterial cell is currently not understood.

Type I segregation systems can be subdivided into two families, Ia and Ib, based on the ParA sequence ^23^, with Type Ia ParA proteins possessing an additional N-terminal helix-turn-helix domain (NTD) that is involved in site-specific DNA binding for *par* gene transcription repression^14,24^. Type Ia systems are found in low-copy number plasmids, whereas Type Ib systems are predominantly present in bacterial chromosomes. The organization of the locus also differs between these two subtypes: In type Ia, the *parA* gene is located directly after the promoter sequence, followed by *parB* and *par*S. In contrast, in type Ib systems, *par*S is located after the promoter, followed by *parA* and *parB*.

The ParA superfamily is highly divergent, with low sequence identity between homologues. Nonetheless, core conserved regions are vital amongst ParAs, particularly the dimerization and nucleotide binding and DNA binding sites^25^. The crystal structure of ParA has been solved in a range of bacterial species and plasmids^5,26–28^, which revealed that despite low sequence identity the overall structure is conserved, and that they form dimers along the same interface. Negative-stain electron microscopy of several ParA orthologues, both of the Type Ia and type Ib families, have shown the formation of filaments in the presence of nucleotide and/or DNA^17,27–29^; However, crystal structures of ParA proteins bound to DNA did not provide any support for filamentous architecture^27^. Whether ParA proteins form filaments, and the molecular basis for filament assembly, remains controversial.

*Vibrio cholerae is* a gram-negative bacterium, and the aetiological agent of cholera, a severe diarrheal disease affecting an estimated 3-5 million worldwide^30^. *V.cholerae* possesses two chromosomes: chromosome 1 (Chr1) and chromosome 2 (Chr2), that are ~3Kbp and ~1Kbp, respectively^31^. Each chromosome encodes its own segregation complex, with Chr1 having encoding a chromosomal Type Ib system, and Chr2 encoding a plasmid-like Type Ia system (Figure 1a). During cell division, both chromosomes segregate in synchronization. Chr1 initiates segregation first, from the old cell pole to new in an asymmetric manner. Once Chr1 reaches the mid-cell region, Chr2 commences segregation. Chr2 segregates symmetrically moving from the mid-cell to quarter cell positions, both chromosomes terminating segregation in unison^32–34^.

**Figure 1:**
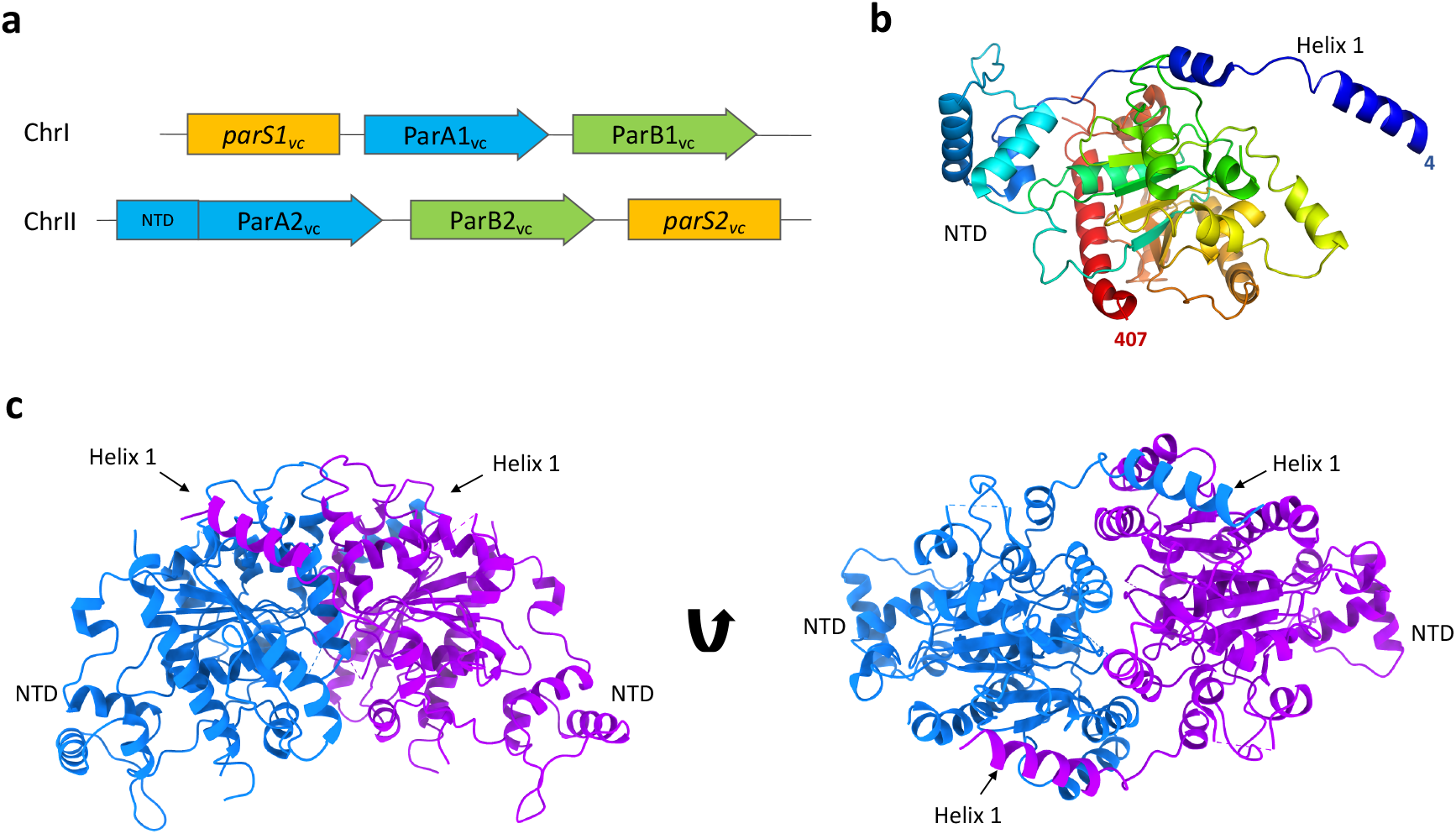
Structure of the ParA2_vc_ dimer. **(a)** Schematic representation of the chromosome segregation systems for the *V. cholerae* chromosome 1 (top), and chromosome II (bottom). The centromere like partition site is in yellow *(parS*), the adapter *parS* binding protein is in green (ParB) and the motor protein is in blue (ParA), with the additional N-terminal domain (NTD) found only in ChrII shown. **(b)** Cartoon representation of the ParA2_vc_ crystal structure, in rainbow coloring, starting from blue at the N terminus, to red at the C terminus. **(c)** The crystallographic dimer of ParA2_vc_ shown from the side (left) and top (right), with the two symmetry-related pairs in blue and magenta, respectively. The NTD and helix 1 are indicated.

Recent studies on the *V. cholerae* Chromosome 2 ParA (ParA2_vc_) have shown that it is a weak ATPase, and binds non-specifically to DNA, similar to other ParA orthologues^29,35^. It also revealed that it forms higher-order assemblies in the presence of DNA, with negative-stain EM analysis confirming the formation of filaments. ParA2_vc_ binds ATP, leading to a slow conformational change to a DNA-binding active state. This then licenses ParA2-ATP dimers to cooperatively bind onto DNA to form higher order complexes. ParB2_vc_ stimulates ParA2_vc_’s ATPase activity, leading to its dissociation from DNA^35^. This fast ParA2 disassembly from the partition complex, coupled with rate-limiting nucleotide exchange, was proposed to generate dynamic ParA2 gradients in *V. cholerae* cells. However, it is not known how ParA2 forms higher order assemblies on DNA, what are the structural changes of ParA2 dimers upon DNA binding and the role of these filaments in ParA2 dynamic gradients.

Here, we report the structure of the ParA2_vc_ filament bound to DNA, determined by cryo-EM. This structure reveals an unexpected set of contacts along the length of the dimer, and onto the DNA. We also have determined the crystal structures of ParA2_vc_ in the apo and nucleotide-bound states. Collectively, these structures reveal a remarkable remodelling of the ParA2_vc_ dimer upon filament formation, providing a structural basis for the cooperativity of its DNA binding, and suggest a molecular mechanism for type I segregation systems.

## Results

### Crystal structure of ParA2_vc_

ParA2_vc_ has low sequence similarity other ParA homologues, with the *E.coli P1/*P7 plasmid ParAs being the closest orthologue of known structure (29% identity). We therefore sought to characterize its structure, to verify that it adopts a similar architecture to other ParA proteins, and to identify any differences with other ParA orthologues.

To this end, we purified ParA2_vc_, and used negative-stain EM to image ParA2_vc_ particles, in a range of nucleotide states. As shown on figure S1, 2D classification of these particles shows the presence of a 2-lobed, V-shaped structure, consistent in shape and dimensions to the P1 ParA dimer reported previously^26^. This suggests that ParA2_vc_ also forms dimers, in all the nucleotide states, as well as in the absence of nucleotides, as we also observed by SEC-MALS previously^35^. It is noteworthy that we did not observe any higher order oligomerization/filament formation in any nucleotide state, unlike that reported in some other ParA orthologues^17,18^, but similar to a previous study on ParA2_vc_^29^.

We next sought to determine the structure of the ParA2_vc_ dimer. The purified protein crystallized readily, and the obtained crystals diffracted to ~ 2.5 Å. We were able to solve the structure by molecular replacement using the P7 ParA crystal structure as a template (see Materials and Methods for details), which allowed us to build an atomic model (Table 1).

**Table 1.**
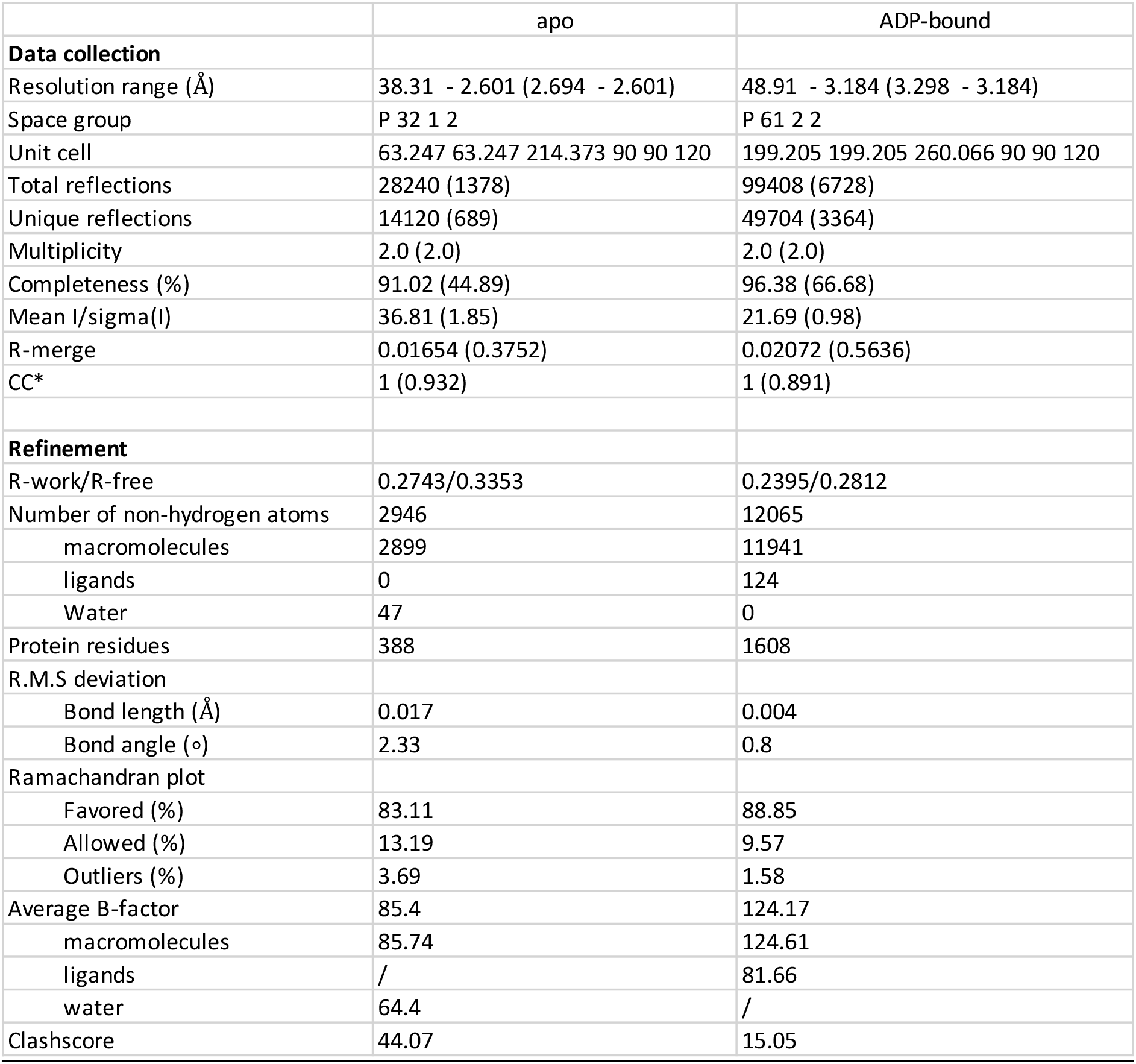
Crystallography and refinement statistics.

The obtained ParA2_vc_ crystal structure includes one ParA2_vc_ molecule per asymmetric unit (Figure 1b). The overall structure of the ParA2_vc_ monomer is similar to that of P7 ParA, with an overall RMSD of 2.2 Å for CA atoms. In particular, the N-terminal HTH domain resembles that of P1 and P7 ParA, confirming that ParA2_vc_ belongs to the type Ia family (Figure 1b, S2). We do nonetheless note some significant differences with these structures, most significantly in the position of the N-terminal helix (helix 1), present in both orthologues, but whose position differs significantly (Figure S2). No ligand density was observed in the active site, confirming that this structure corresponds to the apo state of the protein. It is worth noting that the nucleotide binding site is generally poorly resolved, and in particular we were not able to build the P-loop in this structure.

As indicated above, our EM analysis suggests that ParA2_vc_ is able to form dimers in the absence of nucleotide, at least at high concentration. We therefore questioned if crystallographic symmetry-related ParA2_vc_ molecule pairs might recapitulate the biological dimer. As shown on Figure 1c, one of the symmetry-related pairs is consistent with a biological dimer, and largely resembles the P7 ParA dimer structure (figure S2). This suggests that we have obtained the structure of the ParA2_vc_ dimer through crystallographic symmetry. Comparing the ParA2_vc_ dimer to that of the P7 ParA structure (Figure S2) confirms that they possess a similar architecture, with helix 1 forming a domain-swapped interaction with the adjacent subunit. As indicated above, there is a difference in the position of helix 1, and as a consequence, the relative orientation of the dimer is slightly shifted in the ParA2_vc_ dimer compared to the T7 ParA dimer (Figure S2).

Overall, this structure confirms the common architecture of ParA proteins, and is consistent with the fact that the *V. cholerae* chromosome 2 is plasmid-like, as suggested previously^36^, since the ParA2_vc_ structure confirms that it belongs to the Type Ia family.

### Structure of ParA2_vc_ bound to nucleotide

Previous studies on the Type Ia P1/P7 ParA suggested structural rearrangements of the ParA dimer upon binding to ATP^26^. We therefore sought to identify if the ParA2_vc_ structure was altered in the presence of nucleotide. To address this, we performed crystallization trials in the presence of ADP, ATP, and the non-hydrolyzable ATP analogue ATPγS. While crystals grew and diffracted in all co-crystallization experiments, in most cases the crystals possessed the same crystal form as the apo structure described above, and no nucleotide was observed in the active site for these. Nonetheless, one dataset collected on a crystal obtained in the presence of ADP showed a different space group (Table 1). Molecular replacement was performed using the apo ParA2_vc_ structure, and revealed four molecules per asymmetric unit, consisting of two ParA2_vc_ dimers (Figure 2a, S3a).

**Figure 2:**
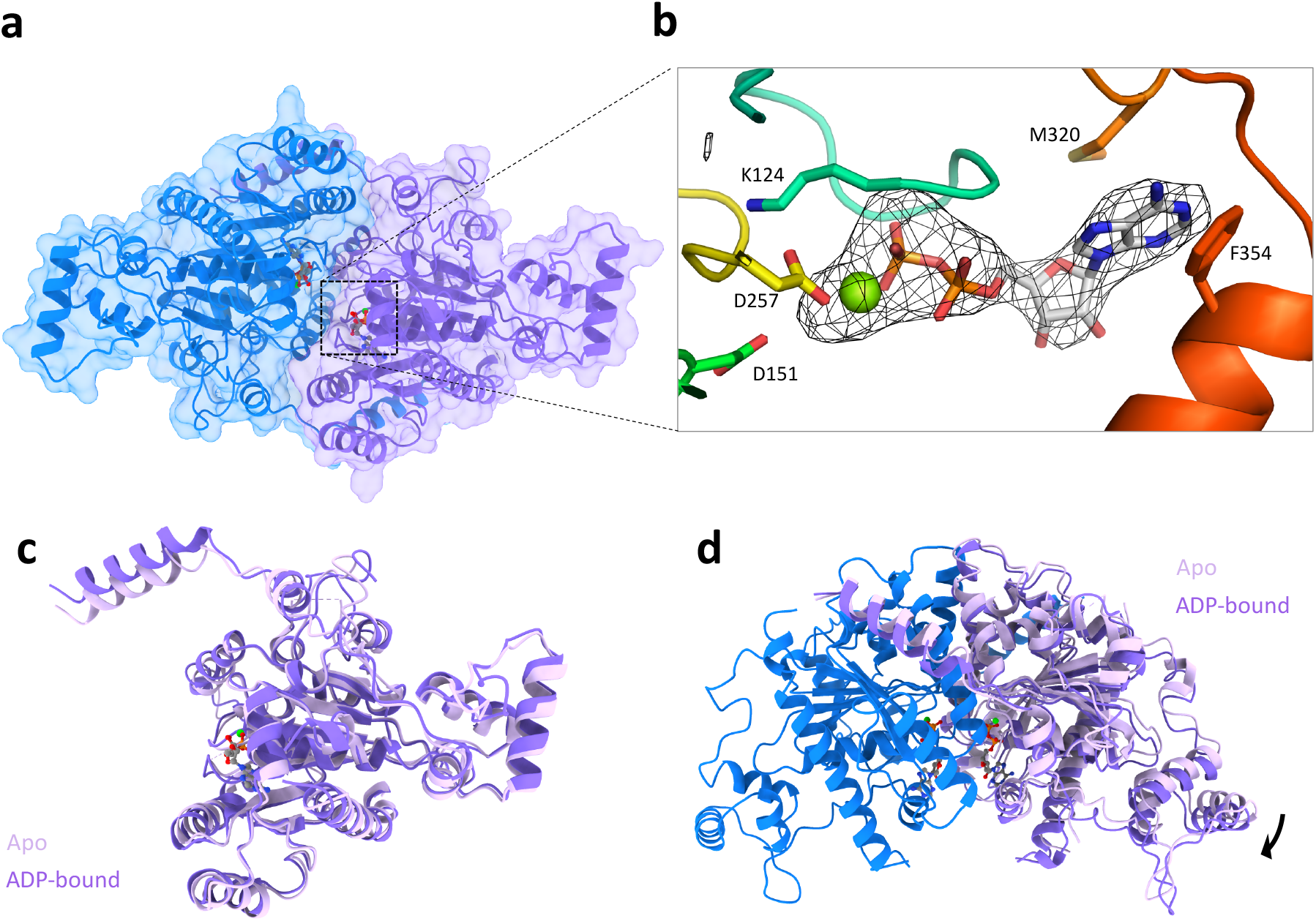
Structure of ParA2_vc_ bound to ADP. **(a)** Surface representation of the crystal structure of a ParA2_vc_-ADP dimer, coloured as in figure 1c. The ADP and Mg molecules, present in each ParA2_vc_ molecules are shown in sticks and sphere representations, respectively. **(b)** Close-up view of the ADP and Mg in one of the ParA2 molecules. The composite omit map is shown around the nucleotide. Residues that interact with the nucleotide and Mg are shown in sticks. **(c)** and **(d)** Overlay of the ParA2vc structure in the apo state (light pink) and ADP-bound state (dark pink). A monomer of each structure is shown in (C), and the dimer is shown in (d), overlaid on the blue subunit; only one chain of the apo structure is shown for clarity.

There is very little difference (RMSD ~ 0.3 Å) between the two ParA2_vc_ dimers contained in the ASU (Figure S3b), as well as in the relative subunit orientation between both dimers (Figure S3c). It is also worth mentioning that the quality of the electron density map is significantly better in the ADP-bound structure compared to the apo structure, especially in the nucleotide binding site, despite its lower resolution (See Materials and Methods for details). We speculate that this might reflect that the nucleotide stabilizes the ParA2_vc_ dimer, thus leading to improved diffraction data, as observed previously for P1 ParA^13^. This is also in agreement with thermal melting assay data, which suggests a stabilization of ParA2_vc_ in the presence of nucleotide^35^.

Ligand density was observed in the active site of all four molecules (Figure 2b), confirming that this structure corresponds to the ADP-bound state of the protein. As shown on Figure S4, the position of the nucleotide is largely similar to that of other ParA orthologues.

As expected, the overall structure of the ParA2_vc_ monomer in the ADP-bound state is similar to that of apo structure (Figure 2c), with a RMSD of ~ 0.8 Å. We do note some changes in the nucleotide binding site, with the P-loop being better ordered in the ADP-bound conformation. In addition, we observed a change in the positioning of the helix 1, which is closer to that of P1/P7 ParA in the ADP-bound structure. As indicated above, helix 1 forms a domain-swapping interaction with the adjacent molecule in the ParA dimer. Because of the difference in the position of this helix, the architecture of the ParA2_vc_ dimer differs between the ParA2_vc_ apo and ADP-states (figure 2d), with a slight shift in the relative subunit position between the two states.

### ParA2_vc_ forms filaments using DNA as a scaffold, regulated by nucleotide hydrolysis

ParA’s ability to form filaments has been highly controversial (See above). ParA filaments have been observed by negative-stain electron microscopy in the presence of nucleotide^16^ and/or dsDNA^27–29^, but fluorescence imaging in cells did not reveal filament formation in multiple systems, and a previously-reported crystal structures of ParA-DNA did not provide evidence for higher-order assembly^27,37^.

As reported above (Fig S1), we did not observe any ParA2_vc_ filament in the absence of DNA, regardless of nucleotide state. However, in the presence of both DNA and nucleotide, filaments were observed by negative-stain EM (Figure S5). Intriguingly, we did not observe filaments in the absence of nucleotide, contrary to a previous study^29^. We also observed that the ParA2_vc_-DNA filaments could only be obtained at high protein concentration, and that they dissociate at lower protein concentration (Figure S5). In contrast, we were able to obtain stable filaments in the presence of the non-hydrolysable ATP analogue ATPγS, which did not dissociate at lower protein concentration (Figure S5c). These results suggest that the assembly of the ParA2_vc_-DNA filament is controlled by ATP hydrolysis, with ATP binding promoting its assembly, and ATP hydrolysis in turn triggering its disassembly. This mechanism is similar to that of Actin and its bacterial homologues, although Actin-like filaments are generally more stable than the ParA2vc-DNA filament^38,39^.

### Cryo-EM structure of the ParA2_vc_-DNA filament

We next used cryo-EM to determine the structure of the ParA2_vc_-DNA filaments described above. As indicated, the presence of non-hydrolysable ATP analogue was required to obtain stable filaments, suitable for cryo-EM analysis. These filaments readily went into ice (Figure 3a), and 2D classification of a resulting cryo-EM dataset confirmed that they are ordered, with the DNA backbone, nucleotide, and secondary structure elements of the protein easily identifiable (Figure 3b). Using this data, we were able to obtain a map to 4.5 Å resolution (Table 2, figure 3b, S6), by helical reconstruction. We then exploited the crystal structure of Par2_vc_ bound to ADP (see above) to build an atomic model of the ParA2_vc_-DNA filament (Figure 3b, S7, and movie S1; see materials and methods for details).

**Figure 3:**
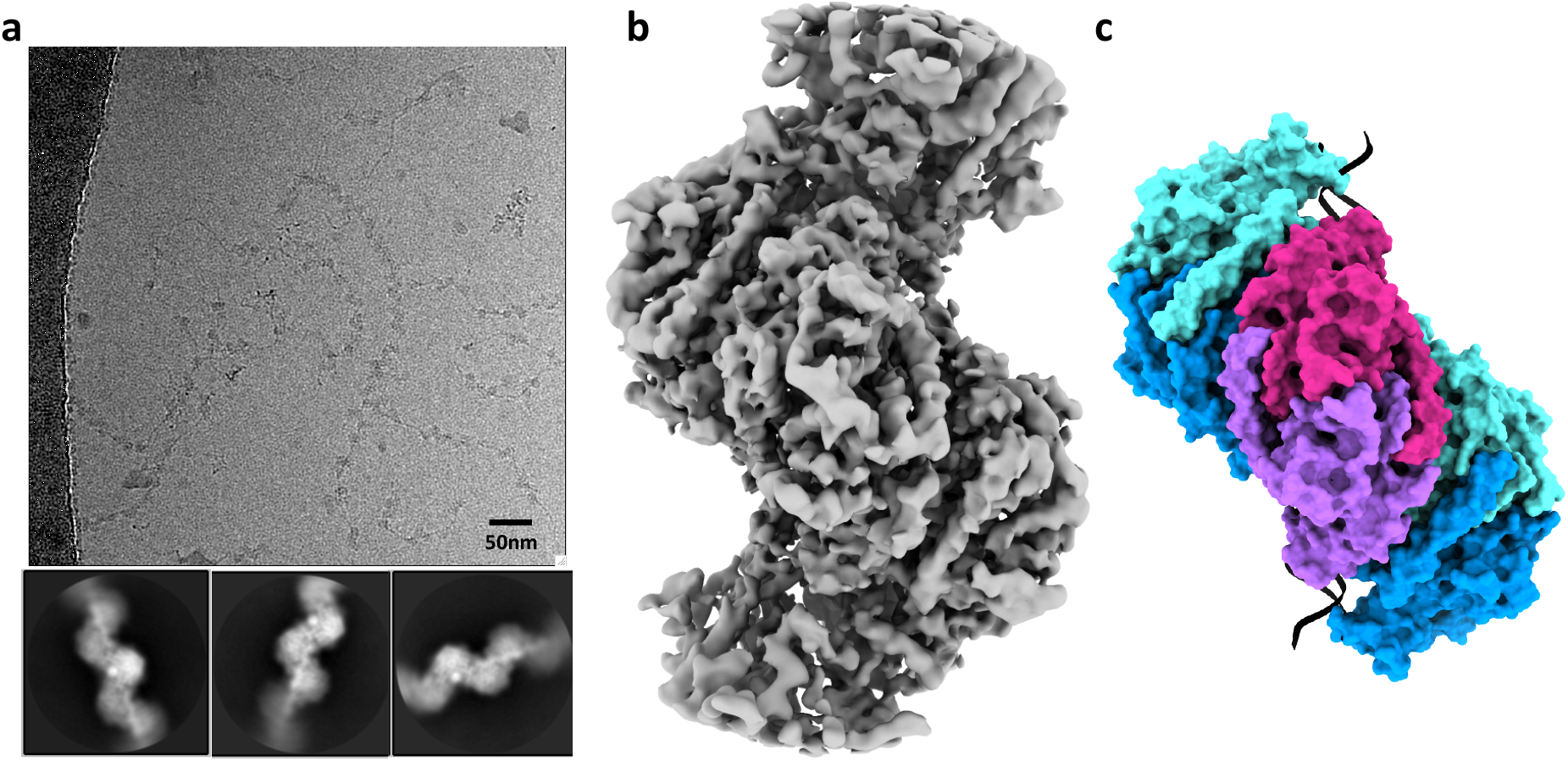
Cryo-EM structure of the ParA2_vc_-ATP_γ_S-DNA filament. **(a)** Representative cryo-electron micrograph of the ParA2_vc_-ATP_γ_S-DNA filament, with selected 2D classes underneath. **(b)** Electron potential map of the ParA2_vc_ATP_γ_S-DNA filament, to 4.5Å resolution. **(c)** Atomic model of the filament structure, covering 3 consecutive ParA2_vc_ dimers bound to DNA in cartoon representation, and in the same orientation as the map in b). Each ParA2_vc_ molecule is colored separately, with the central dimer in purple and magenta, and the two adjacent dimers in cyan and blue, respectively.

**Table 2.** Cryo-EM data collection, processing and model refinement statistics.

|  | ParA2 ATPyS-DNA |
| --- | --- |
| **Data collection and processing** |  |
| Microscope | Titan Krios |
| Magnification | 130,000x |
| Voltage (kV) | 300 |
| Camera | K2 |
| Pixel size (Å) | 1.047 |
| Defocus range (μm) | -2.3 - -1.3 |
| Total dose (e.Å<sup>-2</sup>) | 52.02 |
| Number of micrographs | 5785 |
| Total segments used | 182,997 |
| Symmetry | Helical |
| Rise (Å) | 28.68 |
| Twist (°) | -80.57 |
| Map resolution (Å) | 4.5 |
| **Model refinement** |  |
| Map sharpening B-iso (Å<sup>2</sup>) | 40 |
| Model composition |  |
| Protein residues | 4000 |
| Ligand | 20 |
| Nucleic acid | 116 |
| R.M.S deviation |  |
| Bond length (Å) | 0.008 |
| Bond angle (°) | 1.445 |
| Ramachandran plot |  |
| Favored (%) | 80.48 |
| Allowed (%) | 19.52 |
| Outliers (%) | 0 |
| Average B-factor |  |
| macromolecules | 185.48 |
| ligands | 150.79 |
| Nucleic acid | 129.68 |
| Validation |  |
| Rotamer outliers (%) | 21.02 |
| Clashscore | 39.62 |
| MolProbity score | 3.81 |

As shown on figure 3c, ParA2_vc_ forms a left-handed helix, with a rise of 28.68 Å and a twist of −80.57°. This is consistent with the previously-reported filament architecture, based on low-resolution negative-stain data^29^. The map includes density for five ParA2_vc_ dimers, and a 48bp-long DNA fragment. Density for the DNA is clearly defined (Figure S7), with notably some base pair separation in the best-resolved regions of the map. Density for the ATPγS and Mg molecules is also clearly delineated in the active site (Figure S7).

### ParA2_vc_ interaction with DNA

Our structure of the ParA2_vc_-DNA filament reveals that each ParA2_vc_ molecule binds to DNA via two interaction sites (Figure 4a, movie S2): (1) In the central ParA domain, three regions (residues 322-328, 345-353 and 376-382, figure 4b) interact with the DNA backbone. In particular, a set of basic residues (K326, K327, R350, R352, K376 and K377) form salt bridges with the DNA phosphate. (2) In the N-terminal winged helix-turn-helix domain, the loop between residues 74 and 80 is inserted deep into the minor groove (Figure 4c). Similarly, a number of basic residues (K44, K74, H79) form salt bridges with the DNA backbone. Collectively, these basic residues, mostly present at the positively-charged end of helices, form a continuous positive surface at the bottom of the ParA2_vc_ dimer, ideally suited for interaction with DNA (Figure 4d).

**Figure 4:**
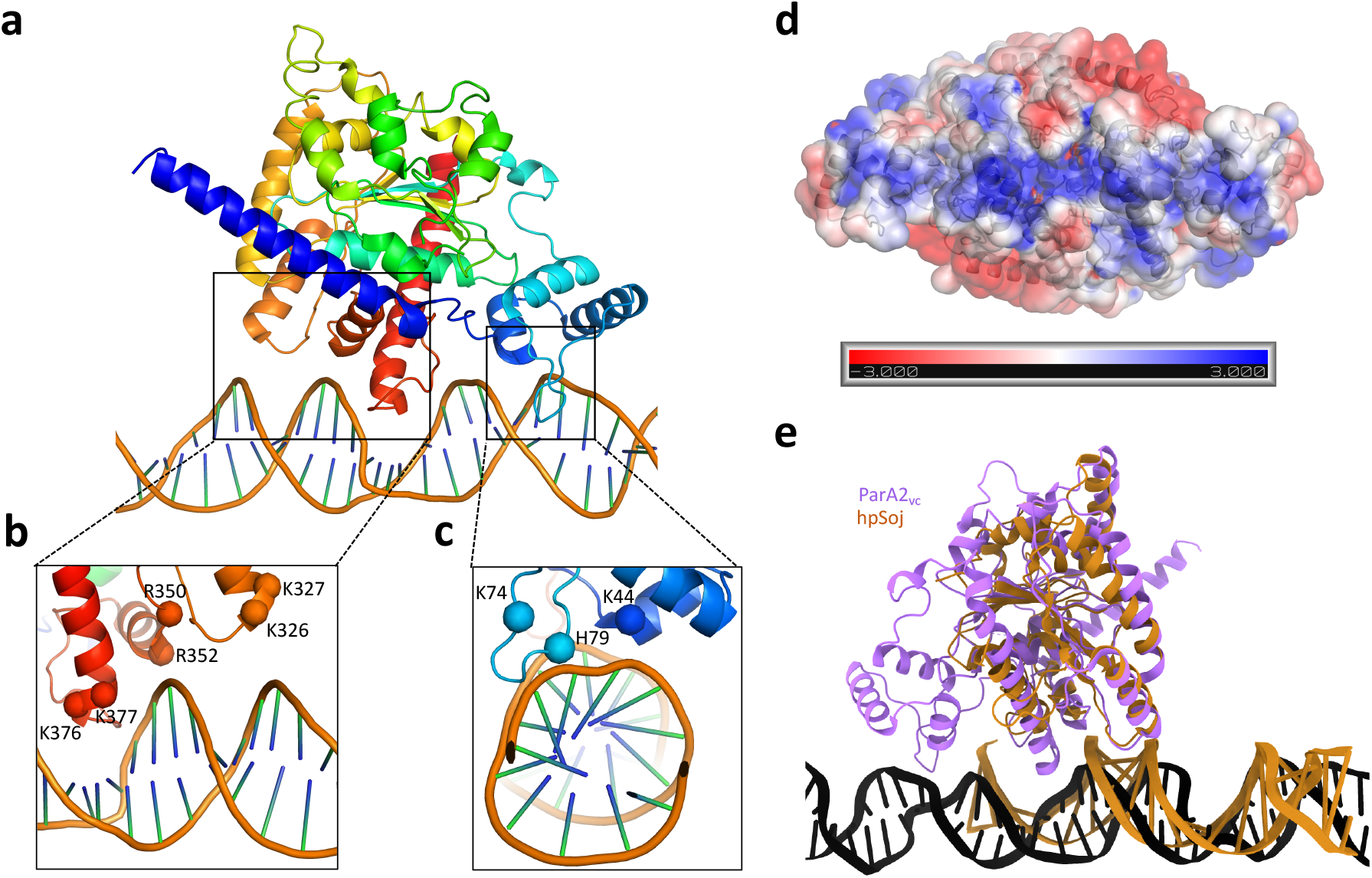
Structural basis for ParA2_vc_’s interaction with DNA. **(a)** A ParA2_vc_ monomer, and the DNA molecule, from the cryo-EM structure are shown in cartoon representation, in rainbow coloring. The two regions forming contacts with the DNA are in black boxes. Closeup views of these two regions are shown in **(b)** and **(c)**, with the basic residues forming salt bridges with the DNA backbone indicated. **(d)** Electrostatic surface representation of the ParA2_vc_ dimer. A positively-charged stretch is clearly present, corresponding to the DNAbinding surface. **(e)** A ParA2_vc_ monomer bound to DNA is shown in purple, overlaid to a _hp_Soj protein bound to DNA in orange. Both proteins interact with DNA on the same surface, but ParA2_vc_ forms additional contacts, via the NTD and the C-terminal helix.

Intriguingly, the basic residues mentioned above are largely not conserved across ParA orthologues, as shown on figure S8. Nonetheless, for two of the corresponding regions (residues 322-328 and 345-353), a number of basic residues are present in all ParA sequences, suggesting a similar mode of binding. Furthermore, co-evolution analysis shows that several of the residues involved in the interaction with DNA, notably K327, R350, R352, K377, have significant evolutionary links with other residues at or near the DNA-binding region Table S1). In keep with this, the recently published structure of a Type Ib ParA protein (the Helicobacter pylori Soj protein, HpSoj) bound to DNA^27^, revealed a largely similar set of interactions with the nucleic acid backbone (Figure 4e). In contrast, the N-terminal domain is not present in Type Ib ParA proteins, and accordingly this set of interactions is not present in the hpSoj-DNA structure. Similarly, while a number of basic residues are found in region 376-382 of type Ia ParA proteins, this region (a loop and part of helix 16) is not found in type Ib ParA proteins, with the exception of the *C. crescentus* ParA orthologue (Figure S8). As shown on figures 4b and 4e, this loop forms a deep insertion within the major groove of the DNA, causing significant distortion of its backbone. As a consequence, the relative orientation of the DNA molecule differs significantly between the hpSoj-DNA crystal structure^27^ and the ParA2_vc_ -DNA structure reported here (Figure S9). Based on the sequence alignment, this difference in DNA orientation can likely be generalized between Type Ia and Type Ib ParA orthologues, although the functional implications are not known. As mentioned above, the *C. crescentus* ParA orthologue (which belongs to the Type Ib family) possesses the additional DNA-binding region near the c-terminus normally found only in Type Ia orthologues, and may therefore possess some common properties between the two families.

It should also be mentioned that the crystal structure of the archaeal plasmid pNOB8 ParA protein^40^ revealed a completely different binding mode to both hpSoj and ParA2vc (Figure S9). In spite of this, the sequence alignment shown on figure S8 indicates that while this protein does not possess the N-terminal domain of Type Ia ParA proteins, all three DNA-binding regions of the core domain are present in the pNOB8 protein, and include several basic residues. It is not known if the difference in DNA interaction corresponds to a crystallization artefact, or reflects biological differences in the interaction with DNA between archaeal and bacterial ParA proteins.

### Filament assembly interface, and remodelling of the ParA dimer upon filament assembly

In the structure of the ParA2_vc_-DNA filament reported here, adjacent ParA dimers form extensive contacts (Figure 5a), with a surface area of ~ 1,500 Å^2^. This interface is largely mediated by three regions: two helices, located at the C-terminus (residues 325-339 and 381-405), and a helix-turn-helix motif from the N-terminal domain (Figure 5b, movie S2), forming electrostatic contacts (Figure S10a). In particular, helices 14 and 16 possess a number of exposed charged residues at the oligomeric surface (Figure S10b), that form salt bridges with the adjacent subunits. We note that it had previously been proposed that only Type Ia ParA orthologues could form filaments, which would be formed only by interactions via the N-terminal domain^29^. However, our structure does not support this, and most of the filament oligomeric interface is located in the C-terminal region of the protein (Figure 5b, S10).

**Figure 5:**
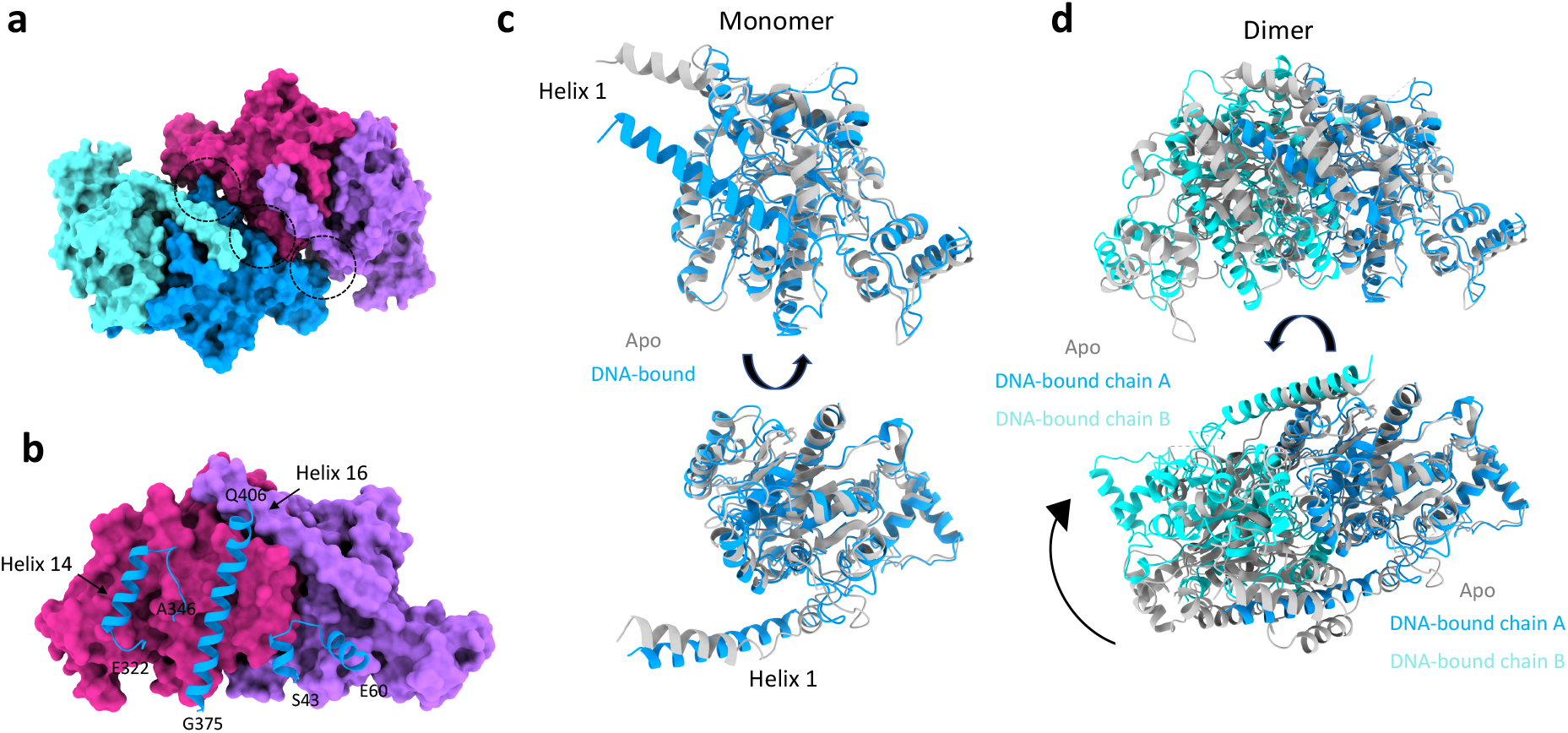
ParA2_vc_ filament interfaces, and structural changes upon filamentation. **(a)** Surface representation of two adjacent ParA2_vc_ dimers in the filament structure, colored as in figure 3c. The interacting regions are indicated with black dotted circles. **(b)** One dimer from (a) is shown in surface representation, and the regions of the second dimer that are involved in the interaction are shown in cartoon. The residue boundaries are indicated. **(c)** and **(d)** Comparison between the ParA2_vc_ structure in the free (grey) and DNA-bound (blue/cyan) conformations. A monomer is shown in (c), illustrating the rearrangement of helix 1; and a dimer is shown in (d), aligned on the blue subunit, to show the dramatic change in the dimer architecture.

Intriguingly, the residues involved in the interface between ParA2_vc_ dimers, within the filament, are not conserved across orthologues, even within the Type Ia family (Figure S8). This could indicate that the filament architecture differs in other bacteria, and/or that some ParA orthologues may not form filaments. Nonetheless, charged residues are found in similar positions in most sequences, and many of these have significant co-evolution links (Table S1). Collectively, this evidence supports the hypothesis that the oligomerization interface is conserved across bacteria and plasmids. Nonetheless, further structural studies of filament architectures in other ParA proteins would be required to verify this.

Finally, another striking feature in the ParA2_vc_-DNA filament structure, is the difference in conformation of ParA molecules, compared to the crystal structures described above. Specifically, helix 1 undergoes a striking conformational change, merging with helix 2 to form a single, extended helix ~ 15 Å from its position in the structures obtained without DNA (Figure 5c). As indicated above, helix 1 forms a cross-dimer interaction, in the ParA2_vc_ dimer. As a consequence, the angle between the two molecules in the filament structure is altered by ~ 30 degrees, compared to the crystal structures described above (Figure 5d, movie S3). This structural rearrangement likely explains the cooperativity in DNA binding, observed in many ParA ortholgues (see discussion), and which have been proposed to be critical for chromosome segregation^41^. Specifically, we propose that the remodeling of the ParA dimer upon DNA binding increases its binding affinity, as an additional binding surface is formed for the binding of additional ParA dimers. We note that residues in helix 1 are not conserved (Figure S8), as observed for the DNA binding and filament interface (see above), but these residues show strong co-evolution links to other residues, closely positioned in the adjacent molecules within the filament structure (table S1), which suggests that the remodelling of helix 1 upon DNA binding is likely applicable to other type Ia ParA orthologues.

## Discussion

In this study, we have reported the structure of the ParA2_vc_ protein, in three states: apo, nucleotide-bound (ADP) and in a filamentous complex with nucleotide and DNA. Importantly, we report the first structure of a ParA protein in the filamentous form. This structure allows us to identify how ParA molecules interact with the DNA, but also how they form higher-order structures. In particular, we show that the NTD forms additional contacts with the DNA, revealing differences between Type Ia and Type Ib ParA proteins. In contrast, we show that the higher-order oligomerization is mostly mediated by the C-terminal region, and co-evolution data suggests that this interface is conserved across ParA orthologues.

From the three structures reported here, we are able to observe the conformational change occurring upon nucleotide binding and filament formation. Combined with prior biochemical and cell-based assays reported previously^29,35^, these structures allow us to propose a mechanistic model for ParA’s higher-order assembly, shown in Figure 6: (a) At physiological concentrations, ParA is at equilibrium between monomeric and dimeric state in the absence of nucleotide. The recruitment of ATP stabilises the dimer. (b) A nucleotide-bound ParA dimer can bind to DNA, and this interaction induces a conformational change to the dimer architecture. (c) This change exposes the filament-forming surface of the DNA-bound ParA dimer, leading to the formation of a filament along the DNA. When encountering a DNA cargo bound to DNA, this activates ParA’s ATPase activity, leading to its disassembly form the DNA, coupled with the release of hydrolysed nucleotide (Figure 6).

**Figure 6:**
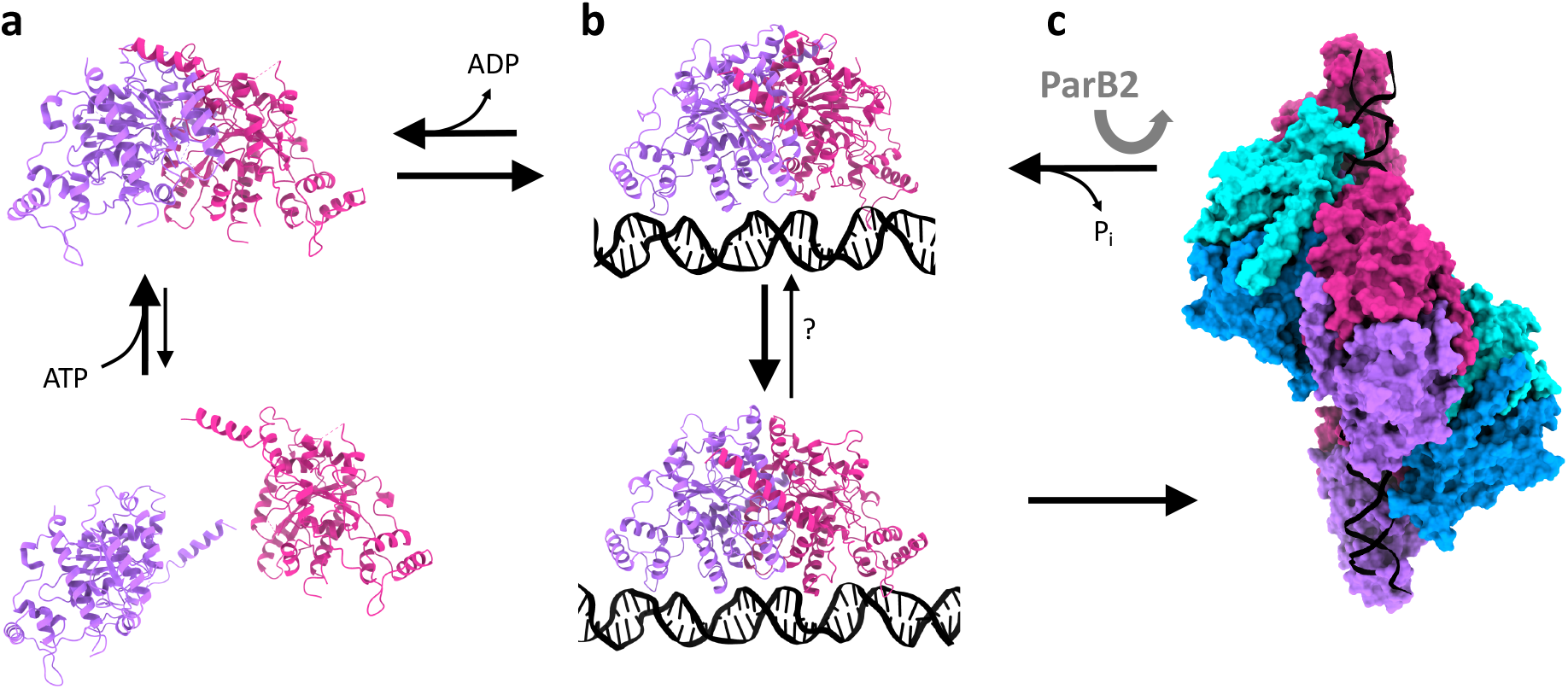
Proposed mechanism for ParA’s cooperative binding to DNA, regulated by ParB. **(a)** in isolation, ParA is in equilibrium between monomer and dimer, with the dimer stabilized by the recruitment of nucleotide. **(b)** In the presence of DNA, the dimer undergoes a dramatic architecture change, exposing its oligomerization interface. **(c)** This leads to the formation of a higher-order assembly, in the form of a short filament segment. The presence of ParB-bound cargo stimulates ParA’s ATPase activity, which leads to its return to the DNA-free conformation of the dimer. This in turn leads to the dissociation of ParA from the DNA, and the release of ADP.

We note that previous biochemical data have shown that ParA2_vc_ binds cooperatively to DNA^35^, as also observed in other ParA orthologues^14,18,28^. The structures reported here likely provide a mechanism for this cooperativity, with the structural changes associated with DNA binding allowing to form a charged surface that permits electrostatic interactions with adjacent dimers. This leads to an increased affinity for the binding of additional ParA2_vc_ molecules adjacent to it. It remains to be verified if the change in dimer architecture is a result of the binding to ATP_g_S, or to DNA.

As mentioned above, whether ParA proteins do form filaments, and the role of such filaments, has remained controversial. In particular, multiple studies using fluorescently-tagged ParA orthologues in dividing cells, revealed that it clusters at high-density chromosomal regions (HDRs)^14,35,42,43^, and do not form filaments across the cell, as required for a mitotic-like mechanism. In keep with this, the negative-stain EM experiments reported here suggests that at near-physiological concentration, the ParA2_vc_ filaments can only form when bound to non-hydrolysed ATP. We therefore propose that *in-situ*, ParA proteins merely form small patches of filaments along the DNA, corresponding to those HDRs observed by fluorescence microscopy. This likely helps forming high-density ParA regions in the nucleoid, as required in the proposed diffusion-ratchet model for segregation.

Nonetheless, a number of questions remain to be addressed to fully validate this mechanistic model. Specifically, while we observed major changes in the dimer architecture from the free ParA to the filament state, it remains to be established if these changes are induced by DNA binding, or by the recruitment of adjacent ParA molecules during filament formation. Furthermore, as indicated above, sequence similarity between ParA orthologue is low, and the residues at the DNA binding regions and filament interface are mostly not conserved. While co-evolution analysis (Table S1) and biochemical studies suggests that these features are likely maintained across species, this remains to be verified experimentally.

It is also worth noting that ParA is structurally similar to MinD, also a P-loop walker type ATPase, but involved in Z-ring localization (Joe Lutkenhaus 2012). MinD forms a dimer, structurally similar to ParA, but does not bind to DNA. Nonetheless, it was recently shown that MinD forms filaments, in the presence of its interacting partner MinC (Andrzej Szewczack-Harris 2019). However, comparison of the MinCD filament to our cryo-EM structure of the ParA2_vc_-ATPγS-DNA filament (Figure S11) reveals that the filaments formed by these two proteins have a completely different architecture, and use different interfaces to form dimer-dimer contacts. Based on this, we postulate that filament formation is not a general feature of this family of proteins, but was adopted independently in ParA and MinD proteins, during evolution.

## Materials and Methods

### Protein expression and purification

For ParA2_vc_ purification, we used the procedure described in Chodha et al.^35^, with a number of modifications. Briefly, cells containing a plasmid including the *parA2_vc_* gene with a T7 promoter, were grown to log phase, expression was induced by adding 1 mM IPTG, the protein was expressed at 16°C overnight, and cells were centrifuged at 6,000g for 15 min. Cell pellets were resuspended in sonication buffer (50mM Tris-HCL pH8, 100mM NaCl, 0.1mM EDTA, 2mM DTT, 50mM (NH_4_)_2_SO_4_) supplemented with 1mg/ml of lysozyme and ½ Roche protease inhibitor tablet (10ml/g of cell pellet). The obtained sample was lysed via sonication for 10 minutes (in 30 s intervals). For electron microscopy experiments, lysed cells were centrifuged at 26,000 × g for 25 minutes, 0.35g/ml of (NH_4_)_2_SO_4_ was added to the supernatant, which was centrifuged as above. The pellet was resuspended in 20ml of Buffer A (50mM Tris pH8, 100mM NaCl, 0.1mM EDTA, 2mM DTT and 20% glycerol), and dialysed against 2 litres of buffer A using 10,000 MWCO SnakeSkin® Dialysis tubing (Thermo Fisher) overnight at 4°C. The sample was then ran through a Heparin HiTrap column (Sigma), and eluted in Buffer A supplemented with 1M NaCl. Fractions containing ParA2_vc_ were then run through a MonoQ column in buffer A and eluted in Buffer A supplemented with 1M NaCl. Finally, the sample was run through a 16/600 pg 200 superdex column (Sigma), in storage buffer (50mM Tris pH8, 500mM NaCl, 0.1mM EDTA, 2mM DTT). Fractions containing ParA2_vc_ were concentrated using a VivaSpin 20 column 10,000 MWCO (Sartorius), before snap freezing in Liquid nitrogen for storage at −80°C. For crystallography, pellets were treated following the method outlined by Chodha et al.^35^

### Crystallization, X-ray crystallography, and structure refinement

ParA2_vc_ at 20mg/ml was set-up for crystallisation trials in sitting drop 96-well plates (Hampton Research), in standard crystallization screens (QIAgen) using a mosquito protein crystallisation robot (sptlabtech), and incubated at 4°C and 20°C for each screen. Crystals were obtained in many conditions, however a significant number of them had a needle-like morphology, and did not diffract. However, we were able to obtain crystals in 0.1M tri-Sodium citrate pH5.5, 20% w/v PEG 3000, grown at 20°C, with a different morphology, which diffracted to ~ 2.5 Å. A dataset was collected for these at the Diamond Light Source beamline io3. The diffraction data was processed to 2.6 Å, and was indexed to the space group P3_2_ 1 2. Phases were obtained by molecular replacement with Phaser^44^ implemented in the Phenix package, using a homology model of ParA2_vc_ based on the P7 ParA structure (3ezF), and missing of the N-terminal residues 1-107. The N-terminal domain was then manually built into the electron density map using *Coot*^45^. The obtained atomic model was refined through multiple cycles of building and refinement, using phenix.refine^46^, to final Rwork/Rfree values of 27.5%/33.5%, respectively (See table 1). The coordinates have been deposited to the PDB, under the accession number 7NPD.We acknowledge that these refinement statistics are relatively poor, and that the geometry of the obtained model is not ideal for the reported resolution, in spite of many cycles of refinement and re-building. This is likely because of the poor quality of the low-resolution data, leading to a poor-quality map and low restraints for refinement. We emphasize that we collected multiple datasets from similar crystals, which all showed similar defects. We therefore suspect that the low quality of the data is due to the flexibility of the molecule, in the absence of ligands in the active site.

For the nucleotide-bound structure, crystallization trials were set up as above, with the ParA2vc sample co-crystallized with 5 mM ADP, ATP or ATPγs and 5 mM MgCl2. As above, crystals were observed in many conditions, however they mostly belong to the needle-like morphology for which no diffraction was obtained, or to the same space group as above. Several datasets were collected nonetheless, but none showed density for ligands in the active site. However, a different crystal morphology was obtained in 24% w/v PEG 1500 with 20% w/v glycerol at 4°C in the presence of ADP. A dataset for such crystal was collected at the Diamond Light Source beamline io3, and diffracted to ~3 Å. The dataset was processed to 3.2 Å, and revealed a space group of P61 2 2. Molecular replacement was carried out using the structure obtained from the data set, as described above, and the structure was refined as described above, to final Rwork/Rfree values of 23.4%/28.1%, respectively (Table 1). The coordinates have been deposited to the PDB, under the accession number 7NPE.

### Negative-stain EM

For the ParA2_vc_ samples, 1mg/ml of protein was incubated with 3.5 mM nucleotide and MgCl_2_ at 30°C for 15 minutes. 1/100 dilutions were made for each nucleotide state into sample buffer containing 100mM NaCl, 50mM Tris pH7.5, 2.5mM nucleotide and 2.5μM MgCl_2_ before staining in 0.75% uranyl formate on glow discharged carbon-coated copper grids (Agar scientific), as described previously.

For the ParA2_vc_-ATPγS-DNA filaments, the samples were prepared as above, except with 2mg/ml of ParA2_vc_. Each sample was then spiked with sonicated salmon sperm DNA (sssNA) (length 1kb) to a final concentration of 0.2mg/ml and left at 30°C for a further 10 minutes, leaving a ParA2_vc_:DNA ratio of ~6:1 and nucleotide and MgCl_2_ at 3mM. A 1/10 dilution for each state was made using sample buffer and staining was done as above. For the ATPγS nucleotide state, ParA2_vc_ at 1 mg/ml was incubated with MgCl_2_ and ATPγS at 4.5mM and incubated at 30°C for 15 minutes. The sample was then spiked with sssDNA (1kb) to a final concentration of 0.14mg/ml and incubated again, resulting in a ParA:DNA ratio of ~3.5:1. From this stock a 1/10 dilution was made, and grid preparation was done as above.

All negative-stain data was collected manually at a defocus range of ~−1 μm to −3.5 μm on a CM100 TEM (Phillips) with a MSC 794 camera (Gatan) at 27,500x with a pixel size of 7.2 Å. For 2D classification, the micrographs were processed using cisTEM^47^, and 2D classes were generated using a box size of 24 pixels.

### Cryo-EM data collection, processing, and model building

ParA2_vc_ at 4 mg/ml was incubated with ~ 6 mM ATPγS and MgCl_2_ at 30°C for 15 minutes, and was then spiked with sssDNA to a final concentration of ~0.8mg/ml of DNA, ~1.9 mg/ml of ParA, and ~4 mM of ATPγS and MgCl_2_. The sample then furtherly incubated for 10 minutes at 30°C. Tween 20 was then added to a final concentration of 0.1% before grid preparation, to facilitate incorporation of the filaments into the holes of the carbon grid. 3 μl of this sample was applied to glow discharged 300 mesh Quantifoil R2/2 grid and left for 30 seconds before manually blotting with filter paper. The grid was then loaded into a Leica EM-GP plunge freezer at 80% humidity and 4°C where a further 3 μl was applied for a 10 second incubation followed by a 4 second blot before being plunged into liquid ethane. Micrographs of ParA2_vc_-ATPγS-DNA filaments were recorded on a 300KV Titan Krios FEG microscope with a Gatan K2 Summit detector in counting mode. 5785 movies were recorded with a pixel size of 1.047Å over 50 frames with a total dose of 52.02e^-^/Å^2^, at a defocus range of −2.3 μm to −1.3 μm. Frames were aligned using MotionCor2^48^, and CTF estimation was obtained using CTFFIND4.1^49^. Filaments were then manually picked in Relion^50^, and helical segments were extracted with a box size of 200 pixels, a tube diameter of 140 Å and a helical rise of 30 Å. 2D classification, 3D classification, and 3D refinement was performed in Relion-3^51^, with a tube density used as an initial reference map. For 3D classification, 6 classes were generated over 25 iterations with a mask diameter of 200 Å over a local helical search range of −70° to −90° for twist and 20 Å to 40 Å for rise. A chosen class was then selected and put into 3D refinement following the same parameters, and the resulting map was post-processed in Phenix^52^ (See figure S6 for the detailed pipeline). The local resolution was determined with ResMap^53^.

To build the atomic model, a polyA-polyT dsDNA molecule was generated in Coot, and placed in the map. A ParA2_vc_ dimer from the ADP-bound structure (see above) was fitted into the EM map using ChimeraX^54^, and helix 1 was re-built manually in Coot. Multiple copies of the resulting dimer were then generated, and placed in the corresponding density in ChimeraX. The final model was then subjected to real-space refinement in Phenix^55^. The coordinates have been deposited to the PDB, under the accession number 7NPF, and the map has been deposited to the EMDB, with the accession number 12515.

### Sequence alignment, co-evolution, and structure representation

The ParA orthologue sequences were aligned with ClustalW^56^, and ESPript^57^ was used to generate the alignment figure. The co-evolution analysis was performed using the GREMLIN server^58^. All structural figures and movies were generated using either ChimeraX, or PyMOL^59^.

## Acknowledgements

A.V.P. was recipient of a PhD scholarship from the Global Strategic Alliance at the University of Sheffield. We are grateful to Dr Satpal Chodha for helpful discussion on ParA biochemistry. We acknowledge the University of Sheffield EM facility for assistance with negative-stain EM data collection, and cryo-EM grid screening. X-ray crystallography data for the ParA2_vc_ apo and ADP-bound were collected at the Diamond Light Source (proposal MX24447), and the Cryo-EM data for the ParA2_vc_-DNA structure was collected at eBIC (proposal EM20970).

## Author contributions

A.V.P., L.C.H, and J.R.C.B. conceived the project and designed the experiments. A.V.P. performed the protein expression and purification, crystallization, X-ray crystallography, negative-stain EM, and cryo-EM experiments, processed the crystallography and EM data, and performed the atomic model building and refinements. S.T. aided with the EM screening and optimization, and DM contributed to the cryo-EM data refinement. A.V.P. and J.R.C.B. wrote the manuscript, with contributions from all the authors.

## References

1 Gordon, G. S. & Wright, A. DNA segregation in bacteria. Annu Rev Microbiol 54, 681–708, doi:10.1146/annurev.micro.54.1.681 (2000).

2 Baxter, J. C. & Funnell, B. E. Plasmid Partition Mechanisms. Microbiol Spectr 2, doi:10.1128/microbiolspec.PLAS-0023-2014 (2014).

3 Reyes-Lamothe, R., Nicolas, E. & Sherratt, D. J. Chromosome replication and segregation in bacteria. Annu Rev Genet 46, 121–143, doi:10.1146/annurev-genet-110711-155421 (2012).

4 Brooks, A. C. & Hwang, L. C. Reconstitutions of plasmid partition systems and their mechanisms. Plasmid 91, 37–41, doi:10.1016/j.plasmid.2017.03.004 (2017).

5 Schumacher, M. A. Structural biology of plasmid partition: uncovering the molecular mechanisms of DNA segregation. Biochem J 412, 1–18, doi:10.1042/BJ20080359 (2008).

6 Jalal, A. S. B. & Le, T. B. K. Bacterial chromosome segregation by the ParABS system. Open Biol 10, 200097, doi:10.1098/rsob.200097 (2020).

7 Funnell, B. E. ParB Partition Proteins: Complex Formation and Spreading at Bacterial and Plasmid Centromeres. Front Mol Biosci 3, 44, doi:10.3389/fmolb.2016.00044 (2016).

8 Jalal, A. S., Tran, N. T. & Le, T. B. ParB spreading on DNA requires cytidine triphosphate in vitro. Elife 9, doi:10.7554/eLife.53515 (2020).

9 Soh, Y. M. et al. Self-organization of parS centromeres by the ParB CTP hydrolase. Science 366, 1129–1133, doi:10.1126/science.aay3965 (2019).

10 Osorio-Valeriano, M. et al. ParB-type DNA Segregation Proteins Are CTP-Dependent Molecular Switches. Cell 179, 1512–1524 e1515, doi:10.1016/j.cell.2019.11.015 (2019).

11 Caccamo, M. et al. Genome Segregation by the Venus Flytrap Mechanism: Probing the Interaction Between the ParF ATPase and the ParG Centromere Binding Protein. Front Mol Biosci 7, 108, doi:10.3389/fmolb.2020.00108 (2020).

12 Volante, A. & Alonso, J. C. Molecular Anatomy of ParA-ParA and ParA-ParB Interactions during Plasmid Partitioning. J Biol Chem 290, 18782–18795, doi:10.1074/jbc.M115.649632 (2015).

13 Vecchiarelli, A. G. et al. ATP control of dynamic P1 ParA-DNA interactions: a key role for the nucleoid in plasmid partition. Mol Microbiol 78, 78–91, doi:10.1111/j.1365-2958.2010.07314.x (2010).

14 Baxter, J. C., Waples, W. G. & Funnell, B. E. Nonspecific DNA binding by P1 ParA determines the distribution of plasmid partition and repressor activities. J Biol Chem 295, 17298–17309, doi:10.1074/jbc.RA120.015642 (2020).

15 Ringgaard, S., van Zon, J., Howard, M. & Gerdes, K. Movement and equipositioning of plasmids by ParA filament disassembly. Proc Natl Acad Sci U S A 106, 19369–19374, doi:10.1073/pnas.0908347106 (2009).

16 Fogel, M. A. & Waldor, M. K. A dynamic, mitotic-like mechanism for bacterial chromosome segregation. Genes Dev 20, 3269–3282, doi:10.1101/gad.1496506 (2006).

17 Ptacin, J. L. et al. A spindle-like apparatus guides bacterial chromosome segregation. Nat Cell Biol 12, 791–798, doi:10.1038/ncb2083 (2010).

18 Ebersbach, G. et al. Regular cellular distribution of plasmids by oscillating and filament-forming ParA ATPase of plasmid pB171. Mol Microbiol 61, 1428–1442, doi:10.1111/j.1365-2958.2006.05322.x (2006).

19 Szardenings, F., Guymer, D. & Gerdes, K. ParA ATPases can move and position DNA and subcellular structures. Curr Opin Microbiol 14, 712–718, doi:10.1016/j.mib.2011.09.008 (2011).

20 Hwang, L. C. et al. ParA-mediated plasmid partition driven by protein pattern self-organization. EMBO J 32, 1238–1249, doi:10.1038/emboj.2013.34 (2013).

21 Taylor, J. A. et al. Specific and non-specific interactions of ParB with DNA: implications for chromosome segregation. Nucleic Acids Res 43, 719–731, doi:10.1093/nar/gku1295 (2015).

22 Havey, J. C., Vecchiarelli, A. G. & Funnell, B. E. ATP-regulated interactions between P1 ParA, ParB and non-specific DNA that are stabilized by the plasmid partition site, parS. Nucleic Acids Res 40, 801–812, doi:10.1093/nar/gkr747 (2012).

23 Gerdes, K., Moller-Jensen, J. & Bugge Jensen, R. Plasmid and chromosome partitioning: surprises from phylogeny. Mol Microbiol 37, 455–466, doi:10.1046/j.1365-2958.2000.01975.x (2000).

24 Hayes, F., Radnedge, L., Davis, M. A. & Austin, S. J. The homologous operons for P1 and P7 plasmid partition are autoregulated from dissimilar operator sites. Mol Microbiol 11, 249–260, doi:10.1111/j.1365-2958.1994.tb00305.x (1994).

25 Hester, C. M. & Lutkenhaus, J. Soj (ParA) DNA binding is mediated by conserved arginines and is essential for plasmid segregation. Proc Natl Acad Sci U S A 104, 20326–20331, doi:10.1073/pnas.0705196105 (2007).

26 Dunham, T. D., Xu, W., Funnell, B. E. & Schumacher, M. A. Structural basis for ADP-mediated transcriptional regulation by P1 and P7 ParA. EMBO J 28, 1792–1802, doi:10.1038/emboj.2009.120 (2009).

27 Chu, C. H. et al. Crystal structures of HpSoj-DNA complexes and the nucleoid-adaptor complex formation in chromosome segregation. Nucleic Acids Res 47, 2113–2129, doi:10.1093/nar/gky1251 (2019).

28 Leonard, T. A., Butler, P. J. & Lowe, J. Bacterial chromosome segregation: structure and DNA binding of the Soj dimer--a conserved biological switch. EMBO J 24, 270–282, doi:10.1038/sj.emboj.7600530 (2005).

29 Hui, M. P. et al. ParA2, a Vibrio cholerae chromosome partitioning protein, forms left-handed helical filaments on DNA. Proc Natl Acad Sci U S A 107, 4590–4595, doi:10.1073/pnas.0913060107 (2010).

30 Faruque, S. M., Albert, M. J. & Mekalanos, J. J. Epidemiology, genetics, and ecology of toxigenic Vibrio cholerae. Microbiol Mol Biol Rev 62, 1301–1314 (1998).

31 Heidelberg, J. F. et al. DNA sequence of both chromosomes of the cholera pathogen Vibrio cholerae. Nature 406, 477–483, doi:10.1038/35020000 (2000).

32 Fiebig, A., Keren, K. & Theriot, J. A. Fine-scale time-lapse analysis of the biphasic, dynamic behaviour of the two Vibrio cholerae chromosomes. Mol Microbiol 60, 1164–1178, doi:10.1111/j.1365-2958.2006.05175.x (2006).

33 Fogel, M. A. & Waldor, M. K. Distinct segregation dynamics of the two Vibrio cholerae chromosomes. Mol Microbiol 55, 125–136, doi:10.1111/j.1365-2958.2004.04379.x (2005).

34 Yamaichi, Y., Fogel, M. A., McLeod, S. M., Hui, M. P. & Waldor, M. K. Distinct centromere-like parS sites on the two chromosomes of Vibrio spp. J Bacteriol 189, 5314–5324, doi:10.1128/JB.00416-07 (2007).

35 Chodha, S. S. et al. Kinetic pathway of ATP-induced DNA interactions of ParA2, a protein essential for segregation of <em>Vibrio cholerae</em> chromosome 2. bioRxiv, 2021.2002.2027.433207, doi:10.1101/2021.02.27.433207 (2021).

36 Kirkup, B. C., Jr., Chang, L., Chang, S., Gevers, D. & Polz, M. F. Vibrio chromosomes share common history. BMC Microbiol 10, 137, doi:10.1186/1471-2180-10-137 (2010).

37 Zhang, H. & Schumacher, M. A. Structures of partition protein ParA with nonspecific DNA and ParB effector reveal molecular insights into principles governing Walker-box DNA segregation. Genes Dev 31, 481–492, doi:10.1101/gad.296319.117 (2017).

38 Izore, T. & van den Ent, F. Bacterial Actins. Subcell Biochem 84, 245–266, doi:10.1007/978-3-319-53047-5_8 (2017).

39 Ozyamak, E., Kollman, J. M. & Komeili, A. Bacterial actins and their diversity. Biochemistry 52, 6928–6939, doi:10.1021/bi4010792 (2013).

40 Schumacher, M. A. et al. Structures of archaeal DNA segregation machinery reveal bacterial and eukaryotic linkages. Science 349, 1120–1124, doi:10.1126/science.aaa9046 (2015).

41 Jindal, L. & Emberly, E. Operational Principles for the Dynamics of the In Vitro ParA-ParB System. PLoS Comput Biol 11, e1004651, doi:10.1371/journal.pcbi.1004651 (2015).

42 McLeod, B. N. et al. A three-dimensional ParF meshwork assembles through the nucleoid to mediate plasmid segregation. Nucleic Acids Res 45, 3158–3171, doi:10.1093/nar/gkw1302 (2017).

43 Le Gall, A. et al. Bacterial partition complexes segregate within the volume of the nucleoid. Nat Commun 7, 12107, doi:10.1038/ncomms12107 (2016).

44 McCoy, A. J. et al. Phaser crystallographic software. J Appl Crystallogr 40, 658–674, doi:10.1107/S0021889807021206 (2007).

45 Emsley, P., Lohkamp, B., Scott, W. G. & Cowtan, K. Features and development of Coot. Acta Crystallogr D Biol Crystallogr 66, 486–501, doi:10.1107/S0907444910007493 (2010).

46 Afonine, P. V. et al. Towards automated crystallographic structure refinement with phenix.refine. Acta Crystallogr D Biol Crystallogr 68, 352–367, doi:10.1107/S0907444912001308 (2012).

47 Grant, T., Rohou, A. & Grigorieff, N. cisTEM, user-friendly software for single-particle image processing. Elife 7, doi:10.7554/eLife.35383 (2018).

48 Zheng, S. Q. et al. MotionCor2: anisotropic correction of beam-induced motion for improved cryo-electron microscopy. Nat Methods 14, 331–332, doi:10.1038/nmeth.4193 (2017).

49 Rohou, A. & Grigorieff, N. CTFFIND4: Fast and accurate defocus estimation from electron micrographs. J Struct Biol 192, 216–221, doi:10.1016/j.jsb.2015.08.008 (2015).

50 He, S. & Scheres, S. H. W. Helical reconstruction in RELION. J Struct Biol 198, 163–176, doi:10.1016/j.jsb.2017.02.003 (2017).

51 Zivanov, J. et al. New tools for automated high-resolution cryo-EM structure determination in RELION-3. Elife 7, doi:10.7554/eLife.42166 (2018).

52 Afonine, P. V. et al. New tools for the analysis and validation of cryo-EM maps and atomic models. Acta Crystallogr D Struct Biol 74, 814–840, doi:10.1107/S2059798318009324 (2018).

53 Kucukelbir, A., Sigworth, F. J. & Tagare, H. D. Quantifying the local resolution of cryo-EM density maps. Nat Methods 11, 63–65, doi:10.1038/nmeth.2727 (2014).

54 Goddard, T. D. et al. UCSF ChimeraX: Meeting modern challenges in visualization and analysis. Protein Sci 27, 14–25, doi:10.1002/pro.3235 (2018).

55 Afonine, P. V. et al. Real-space refinement in PHENIX for cryo-EM and crystallography. Acta Crystallogr D Struct Biol 74, 531–544, doi:10.1107/S2059798318006551 (2018).

56 Hung, J. H. & Weng, Z. Sequence Alignment and Homology Search with BLAST and ClustalW. Cold Spring Harb Protoc 2016, doi:10.1101/pdb.prot093088 (2016).

57 Gouet, P., Robert, X. & Courcelle, E. ESPript/ENDscript: Extracting and rendering sequence and 3D information from atomic structures of proteins. Nucleic Acids Res 31, 3320–3323, doi:10.1093/nar/gkg556 (2003).

58 Ovchinnikov, S., Kamisetty, H. & Baker, D. Robust and accurate prediction of residue-residue interactions across protein interfaces using evolutionary information. Elife 3, e02030, doi:10.7554/eLife.02030 (2014).

59 The PyMOL Molecular Graphics System, Version 2.0 Schrödinger, LLC. .

