## Supplementary material for "The cryo-EM structure of the bacterial type I DNA segregation ATPase filament reveals its conformational plasticity upon DNA binding": Figure S

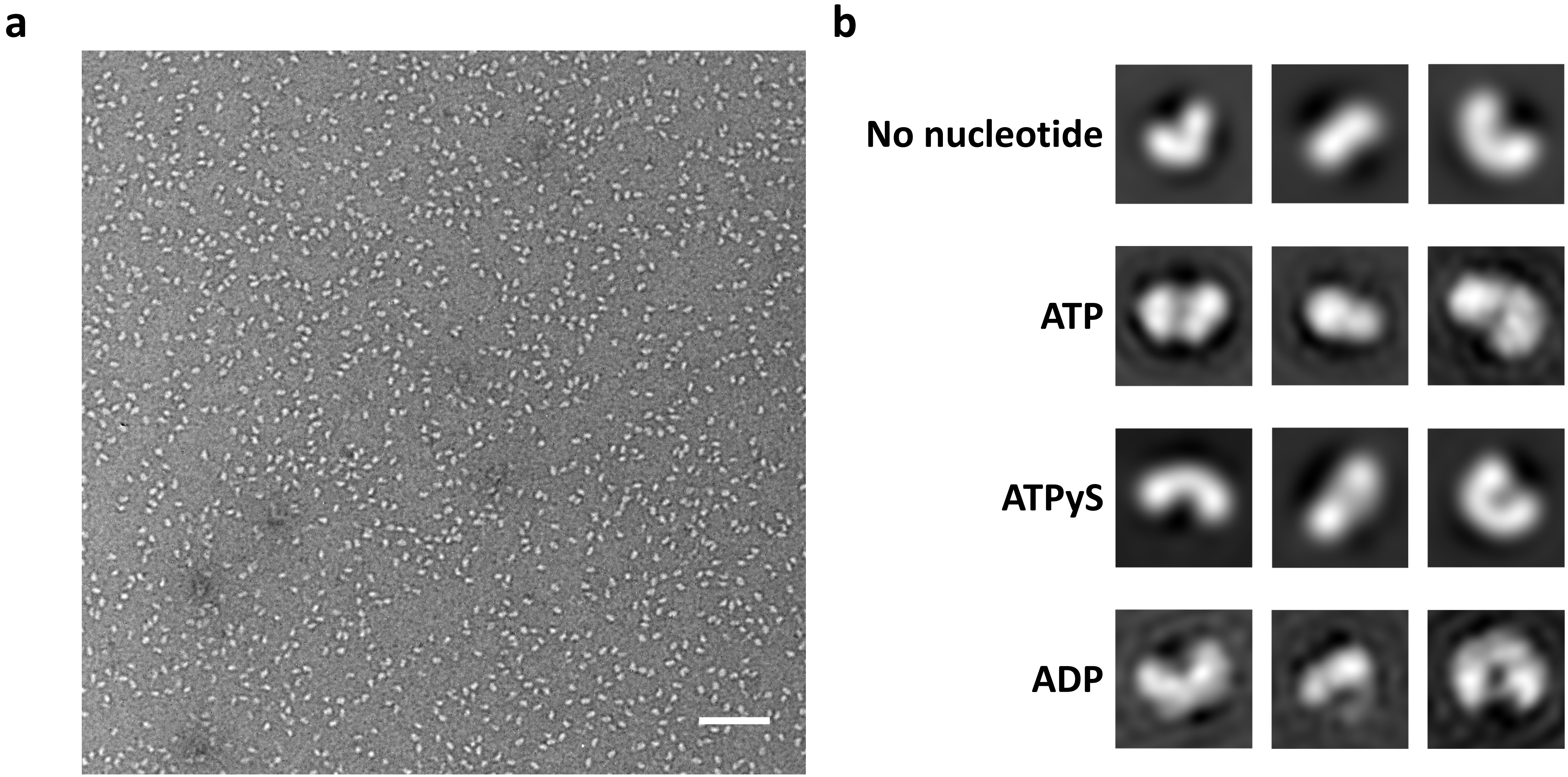


Figure S1: Micrograph and 2D classes of ParA2$_{vc}$ dimers. (a) Negative-stain electron micrograph of ParA2$_{vc}$ in the presence of ATP. Elongated particles, approximately 10 nm long, are visible. The scale bar indicates 100nm. **(b)** 2D classes of particles from negative-stain EM micrographs of ParA2$_{vc}$ in varying nucleotide states. Two-lobbed particles, consistent with a dimer in different orientations, are visible in all states.

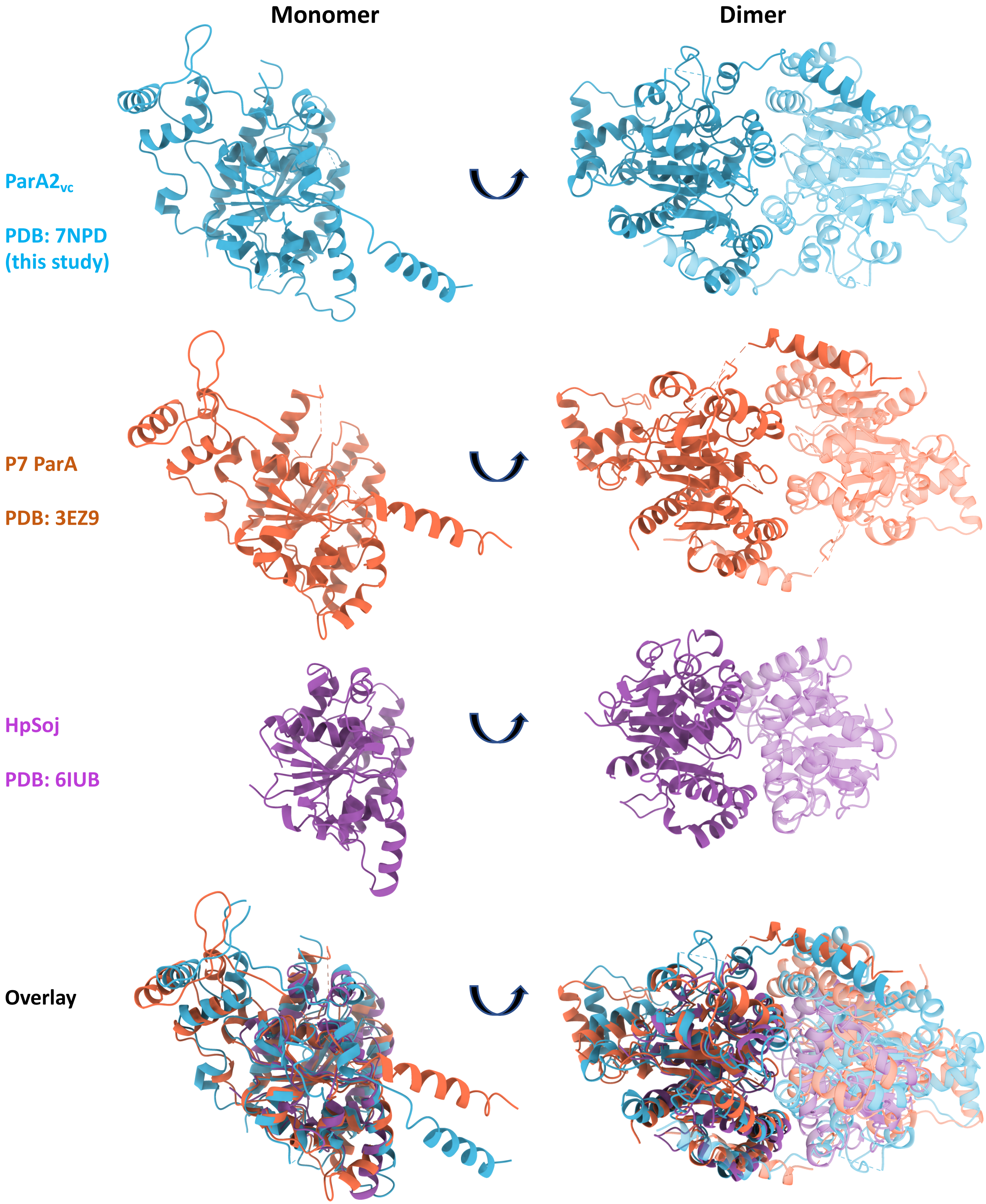


**Figure S2: Comparison of ParA structures across bacterial species.** A monomer (left) and dimer (right) is shown for the structures of $ParA2_{VC}$ (cyan), the *E.coli* P7 plasmid ParA (orange), and $_{Hp}Soj$ (purple). All three structures are aligned at the bottom, on the left chain for the dimer.

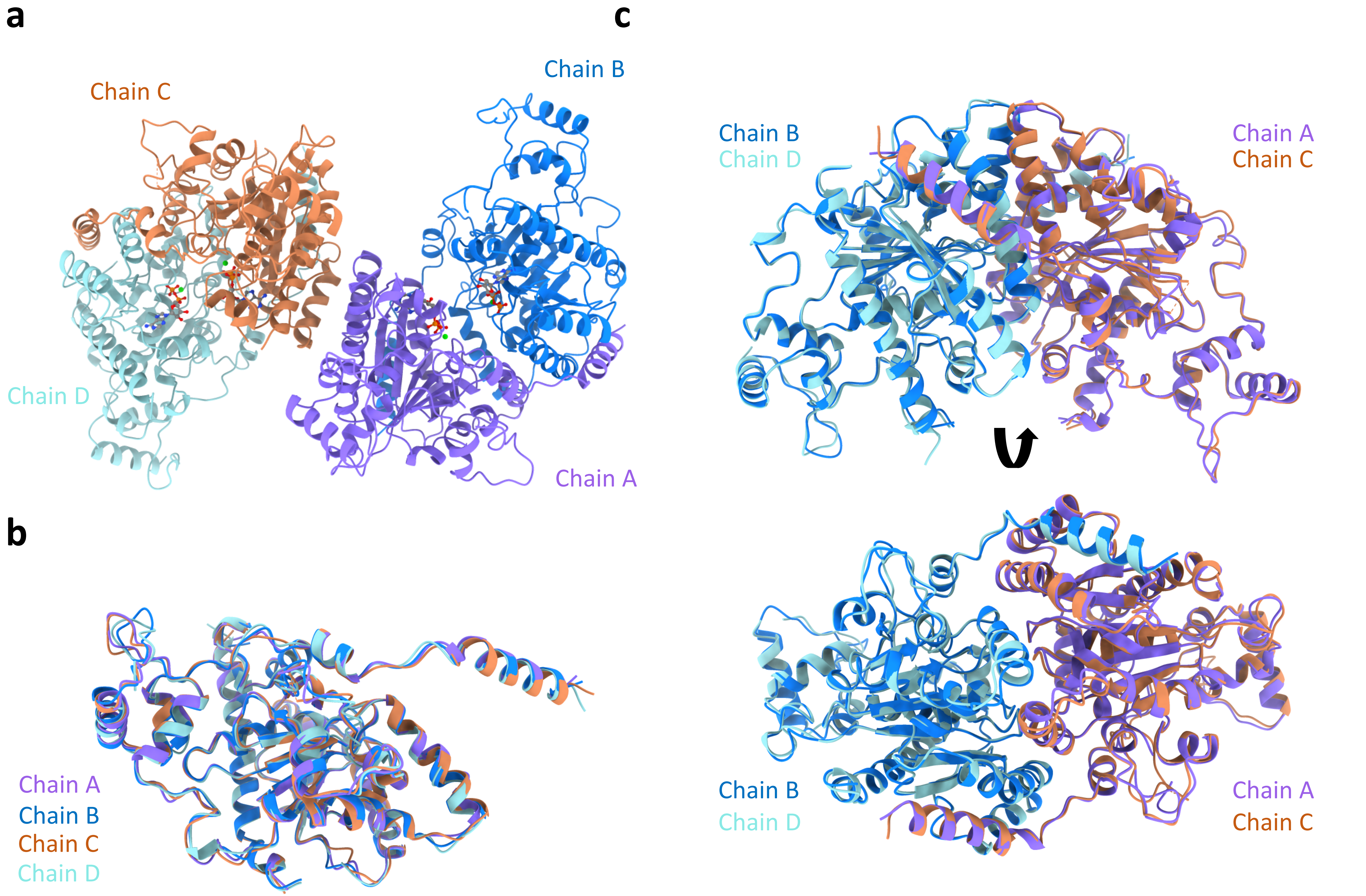


**Figure S3: Comparison of chains in the ParA2$_{vc}$-ADP asymmetric unit. (a)** Cartoon representation of the ParA2vc-ADP structure asymmetric unit, showing 4 protein molecules, in purple, blue, orange, and cyan, respectively. Density for the nucleotide is found in all four. **(b)** Overlay of all 4 chains, coloured as in (a). **(c)** Side (top) and top (bottom) views of the two dimers in the ASU, coloured as in (a). All four chains only show minor variations in loop regions, and the dimeric architecture is the same for the two dimers.

**a**

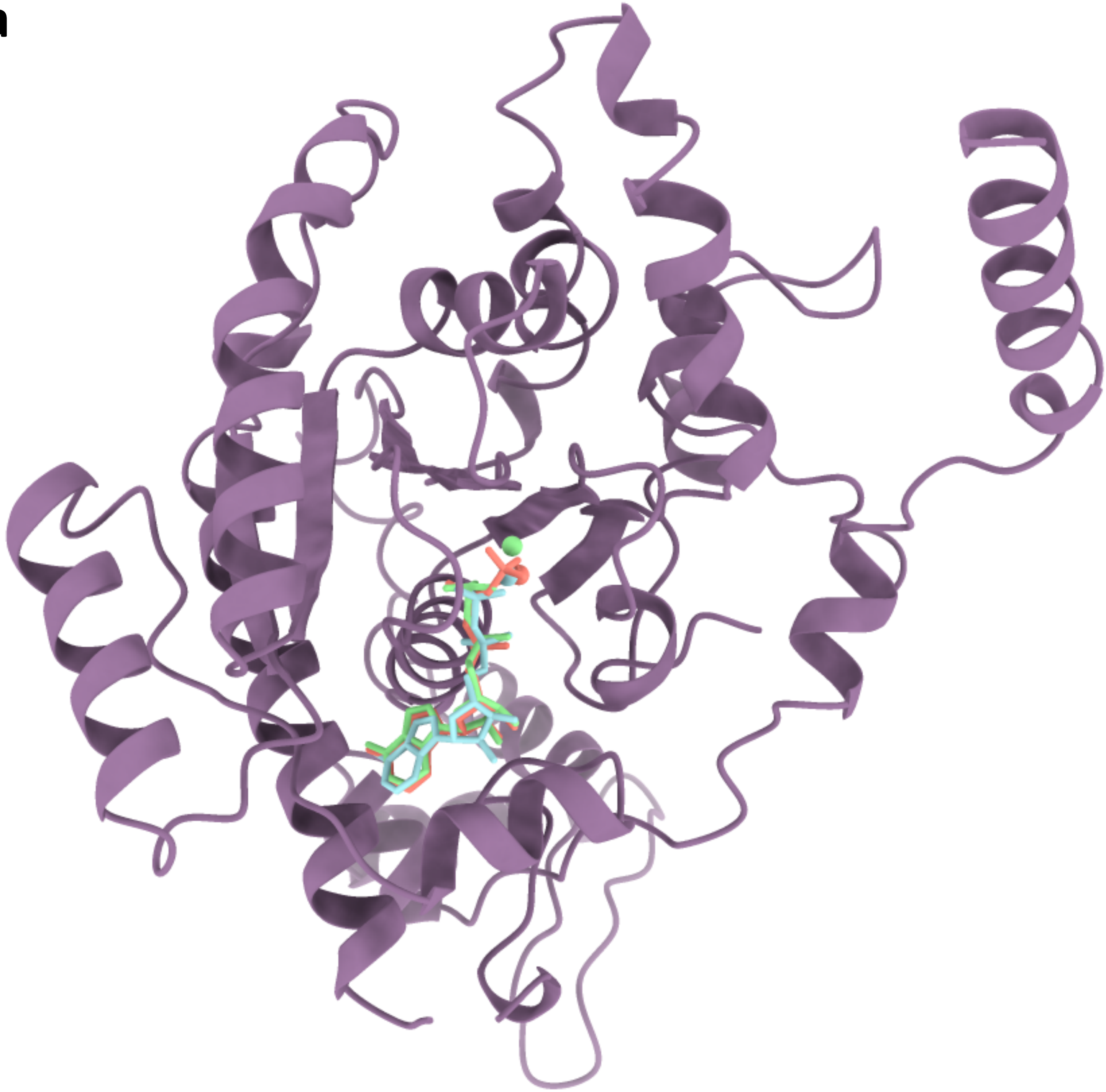

**Figure S4: Nucleotide binding in ParA orthologues.** The structure of ParA2$_{VC}$ is shown in purple, with the bound ADP molecule in cyan. The nucleotide in overlaid P1 ParA structure (ADP:PDB ID 3ZE6) and in hpSoj (ATP:PDB ID 6IUB) are in green and orange, respectively. The nucleotide is positioned similarly in all three structures.

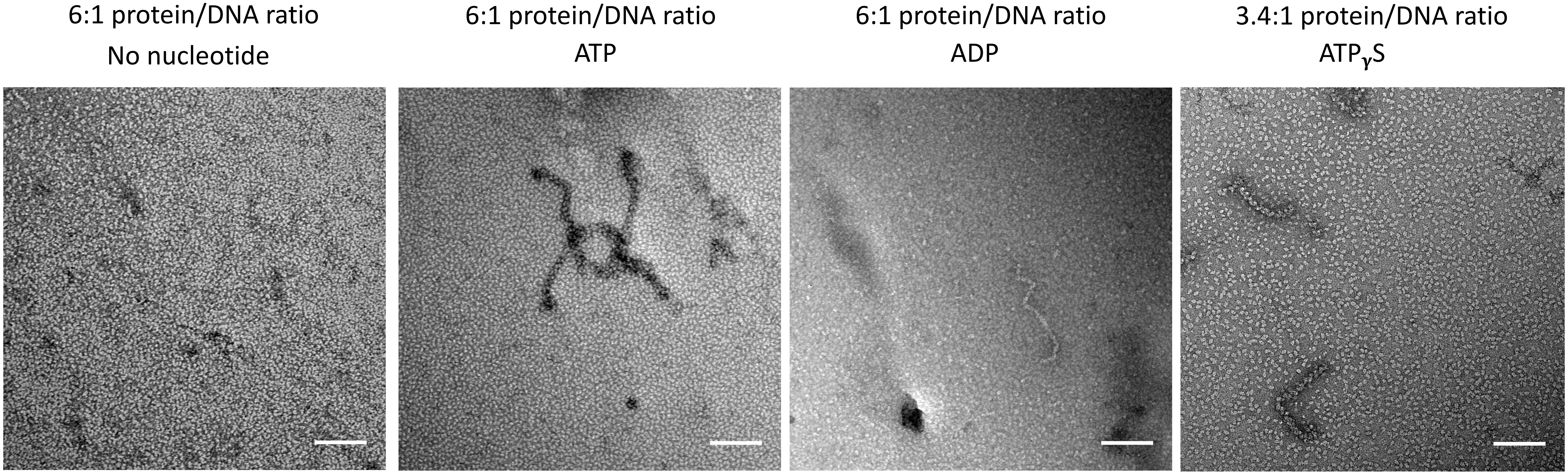


**Figure S5: Negative-stain electron micrographs of $ParA2_{vc}$ with ssDNA, in varying nucleotide states.** The white bar indicates 100 nm. $ParA2_{vc}$ only forms filaments on DNA in the presence of nucleotide, and in large excess of protein. At lower protein concentration, $ATP_{\gamma}S$ is required to observe stable filaments.

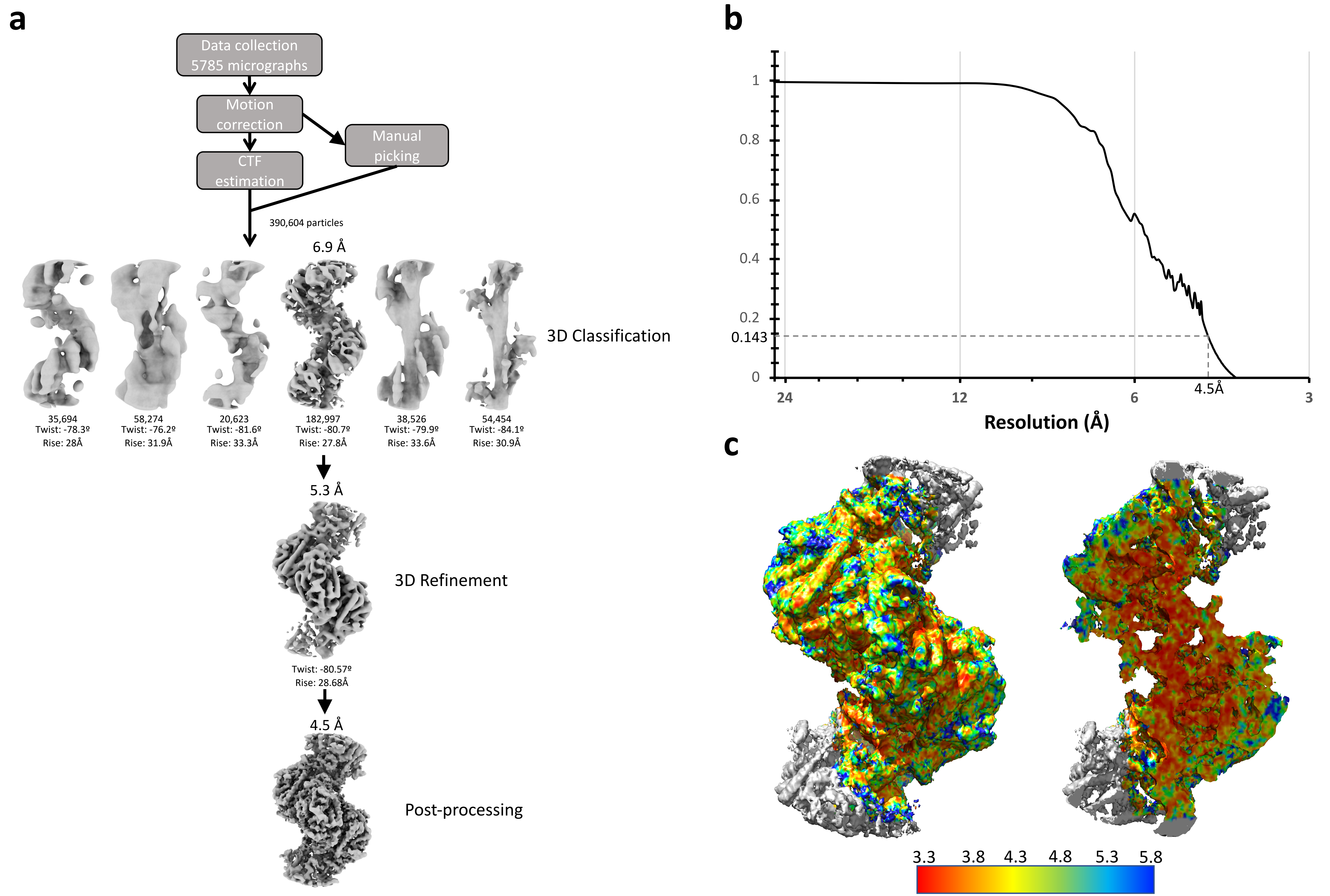


Figure S6: Details of the ParA2$_{vc}$-ATP$_{\gamma}$S-DNA filament cryo-EM structure determination. (a) Data processing pipeline. (b) FSC curve for the masked post-processing refinement. (c) Local resolution of the electron potential map.

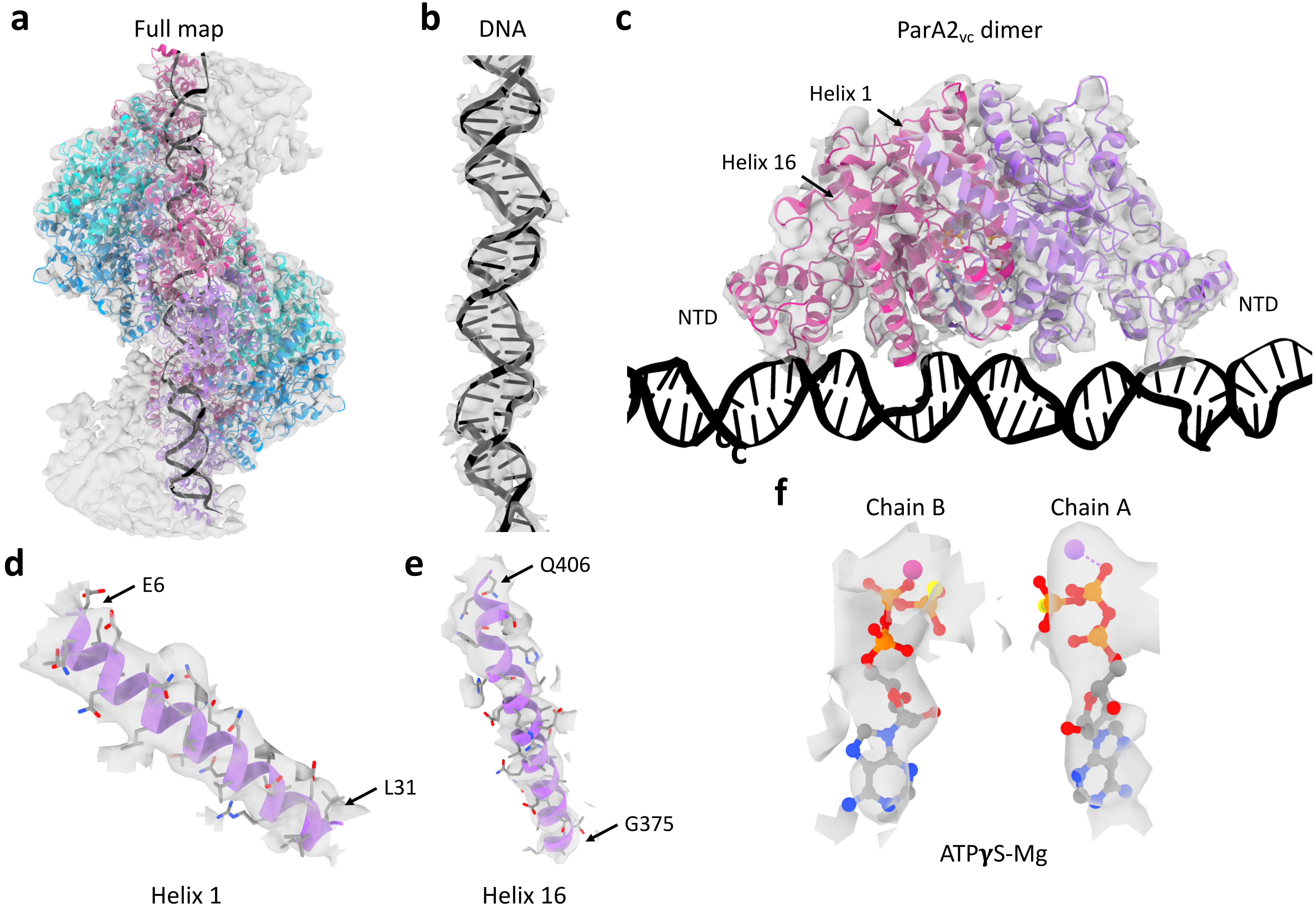


**Figure S7: Examples of the fit of the ParA2$_{vc}$-ATP$_{\gamma}$S-DNA atomic model to the electron potential map.** Various elements are shown, including the full atomic model **(a)**, DNA **(b)**, a single ParA2$_{vc}$ dimer **(c)**, the N-terminal **(d)** and C-terminal **(e)** helices, and the two nucleotides of the central dimer **(f)**. These illustrate that the map features are consistent with the reported resolution.

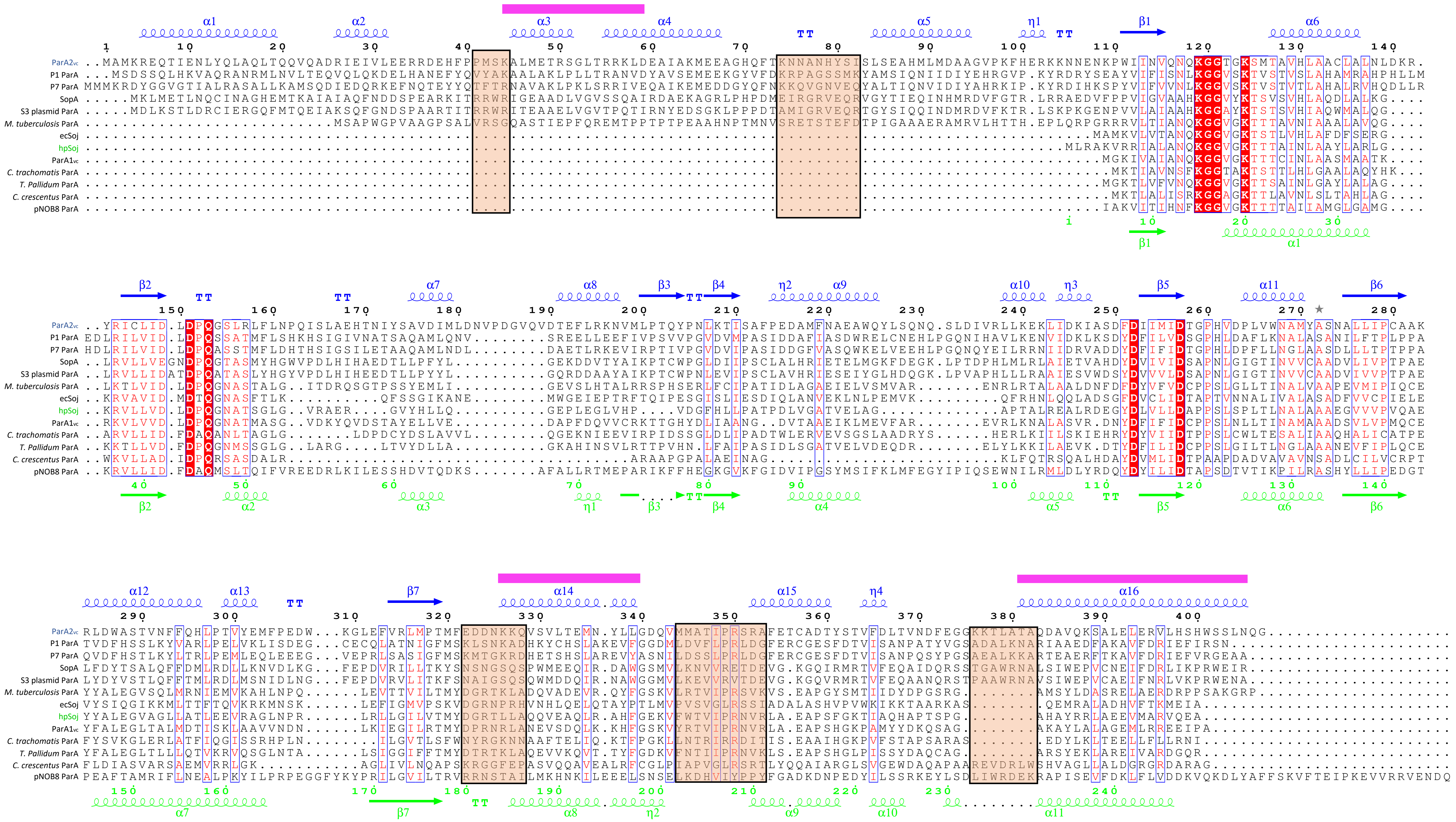


**Figure S8: Multiple sequence alignment of various ParA orthologues.** The first six are Type Ia sequences, the next seven are type Ib sequences, and the bottom sequence is for the archaea pNOB8 plasmid ParA. The secondary structure elements for ParA2$_{VC}$ is in blue at the top, and for HpSoj in green at the bottom. Conserved residues are in red box, similar residues in red characters. The orange boxes indicate regions of ParA2$_{VC}$ that interact with DNA, and the purple bars indicate the filament oligomerization interfaces.

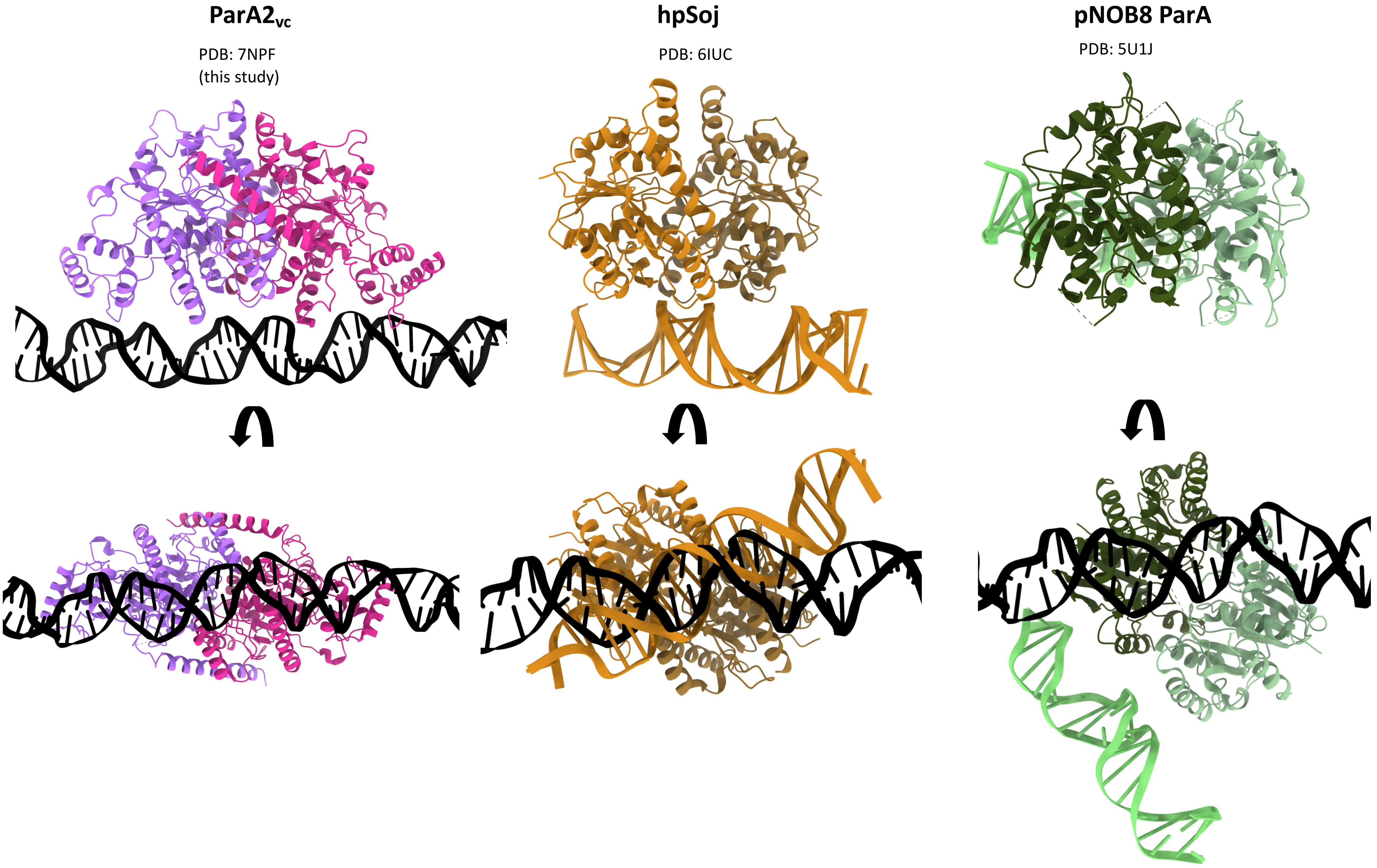


**Figure S9: Comparison of the DNA binding in ParA orthologues.** The structure of the ParA dimer, from the side and bottom, is shown for ParA2$_{vc}$ (left), HpSoj (centre), and pNOB8 ParA (right). The two ParA2$_{vc}$ molecules are colored as in figure 3c, with the DNA in black. For HpSoj and pNOB8 ParA, the position of the DNA in the filament structure reported here is shown in black, for comparison.

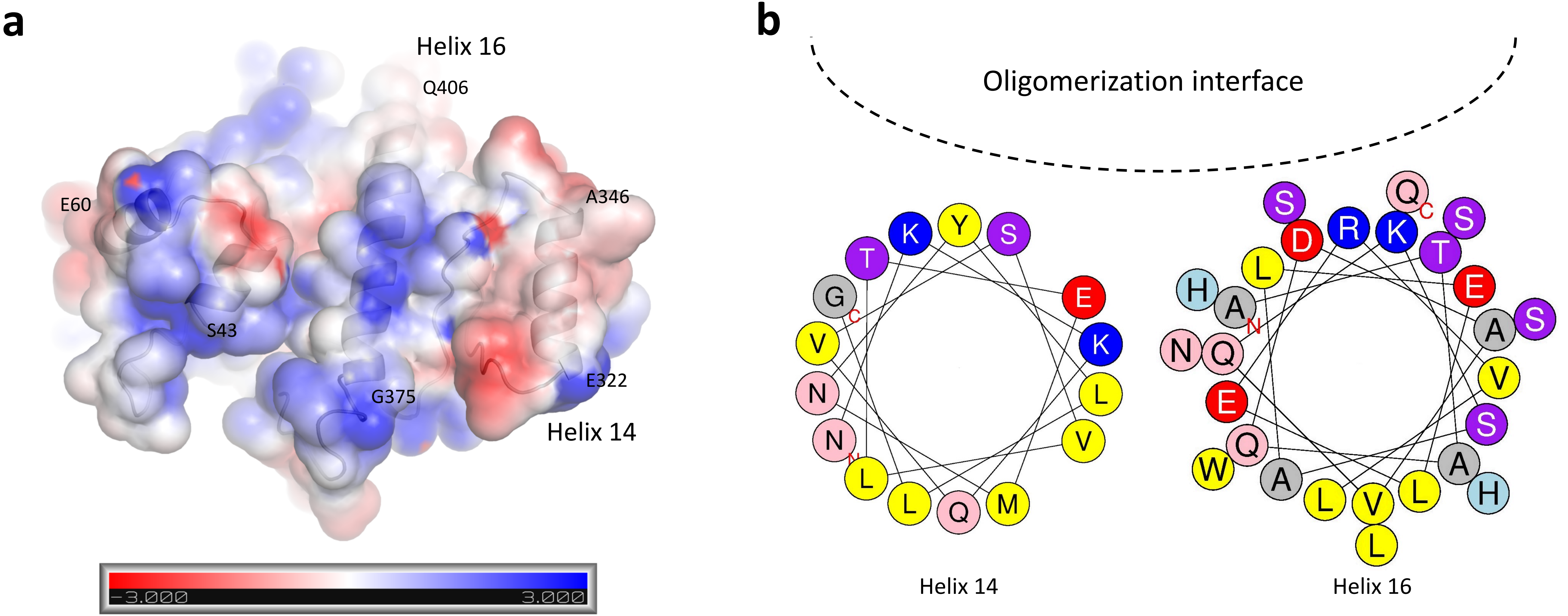


**Figure S10: Molecular details of the oligomerization interface.** **(a)** surface representation of a ParA2$_{VC}$ monomer in the filament structure, colored by charge. The boundaries of the domains involved in dimer-dimer interaction are indicated. **(b)** Helical wheel plot of **α**-helices 14 and 16 of ParA2$_{VC}$, that form the filament oligomerization interface. The charged residues cluster on the interface side.

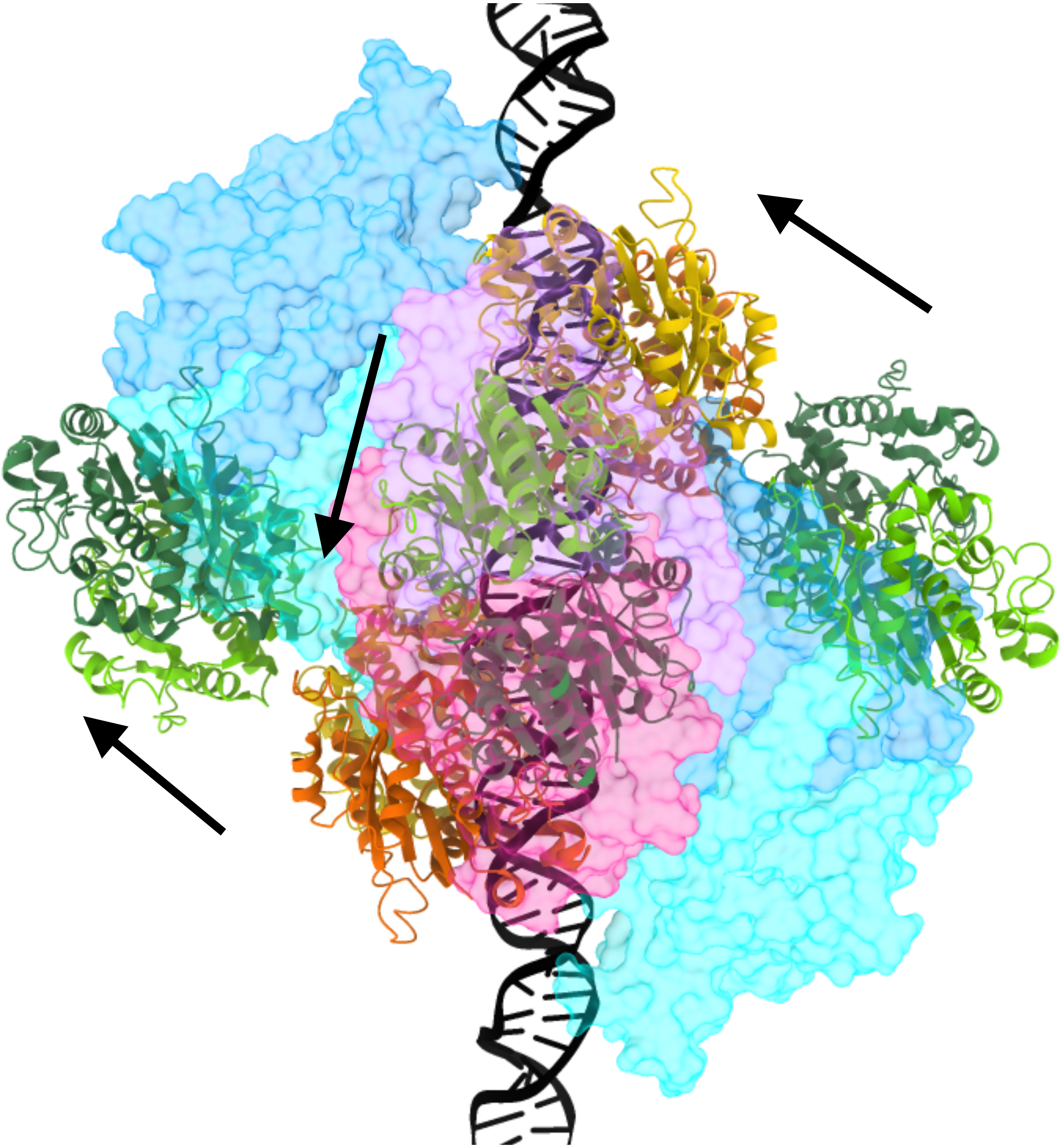

Figure S11. Comparison of the **ParA2$_{vc}$-DNA filament to MinD in the MinCD filament.** A ParA2$_{vc}$ dimer bound to the DNA from the filament structure is shown, represented as in figure 3c. Adjacent MinD dimers from the MinCD filament structure are in light and dark green for one dimer, yellow and orange for the other. The central green MinD dimer is aligned to ParA2$_{vc}$, the arrows indicate the direction of the MinD filament architecture.
